# Pathology of African Swine Fever in wild boar naturally infected with German virus variants

**DOI:** 10.1101/2022.09.14.507889

**Authors:** Julia Sehl-Ewert, Paul Deutschmann, Angele Breithaupt, Sandra Blome

## Abstract

In 2020, African swine fever (ASF) was first notified in German wild boar, reaching a case number of about 4200 to date. Upon experimental infection, pathology is well-documented, however, data on field infections are very scarce in domestic pigs and not available from wild boar, respectively. Although ASF viral genome is considered exceptionally stable, a total of five lineages with 10 distinct virus variants of genotype II have emerged in Eastern Germany. To investigate the pathology in naturally infected wild boar and to evaluate virus variants II, III and IV for their virulence, wild boar carcasses were obtained from three different outbreak areas. The cadavers underwent virological and pathomorphological investigation. Regardless of the virus variant all wild boar revealed characteristic lesions of highest severity indicative for ASF. However, wild boar infected with variant IV from Spree-Neiße (SN) district showed lower viral genome loads and a lower total viral antigen score, but simultaneously revealed more chronic lesions. Our findings indicate a protracted course of the disease at least after infection with variant IV, but need confirmation under standardized experimental conditions. There is a strong need to monitor differences in the virulence among variants to identify potential attenuation that might complicate diagnosis.

## Introduction

Since its first occurrence in Georgia in 2007, African swine fever (ASF) has continuously spread from the Trans-Caucasian region to Russia, and in 2014 further to countries of Europe (1). Last in September 2020, the disease has been confirmed for the first time in a wild boar found in the Spree-Neiße (SN) district in Eastern Germany close to the German-Polish border (2). To date, more than 4200 cases in German wild boar in the Eastern federal states Brandenburg, Saxony and Mecklenburg-Western Pomerania as well as seven outbreaks in domestic pig holdings located in Brandenburg, Mecklenburg-Western Pomerania, Baden-Wurttemberg and Lower Saxony have been officially notified (https://tsis.fli.de/Reports/Info.aspx, visited online on September 9th 2022).

African swine fever, which is caused by the large, enveloped, double-stranded DNA African swine fever virus (ASFV), can occur as acute, subacute, chronic and subclinical disease courses depending on the virulence of the virus strain as well as on the age and immunological background of the animals (3). In European countries except Sardinia, highly virulent virus strains of genotype II are prevalent in domestic and wild pigs typically causing acute-lethal disease similar to a hemorrhagic fever (4–6). Genotype II strains were also identified in the German federal states of Brandenburg and Saxony including the outbreak areas Märkisch-Oderland (MOL), Oder-Spree (LOS), Spree-Neiße (SN) and Görlitz, in which, surprisingly, five lineages (I-V) including a total of ten viral variants (I, II, II.1, III, III.1, IV, IV.1, IV.2, IV.3, V) have emerged due to single nucleotide variations, insertions and deletions affecting different genes including five multigene families (7). More specifically, variants III and IV comprise genetic variations in four multigene family (MGF) genes MGF360-10L, MGF360-15R, MGF100-3L and MGF505-4R while variant II shows variation only in the A240L gene coding for the ASFV thymidylate kinase. Whereas the functions of these genes are largely unknown, ASFV MGF360 and MGF505 have been associated with virulence and pathogenicity of the virus (8, 9). Geographic mapping showed that variant II was predominantly spread in the outbreak area LOS, variant III in MOL, and variant IV in the Southern part of the outbreak area SN as well as in the federal state of Saxony.

To date, macroscopic pathological records of varying depth of detail largely exist only for experimentally ASFV infected domestic pigs (10–16) and less frequently for wild boar (4, 17–20), which is mainly due to the limited access on wild boar and associated difficulties to keep them under experimental conditions. Moreover, histopathological data obtained from animal experiments are much less available, but gained importance in the last years (17, 18, 21). Very recently, the first three reports were published concerning naturally ASFV infected domestic pigs from an outbreak in Vietnam, reporting on the clinical and pathological findings of succumbing and surviving pigs (22, 23) and describing ASF associated age-related lesions (24). In contrast, descriptions of pathological findings of wild boar that succumb to infection under field conditions are completely missing although this animal species is of great relevance in the maintenance and spread of ASFV in Europe. Hence, the diversity and dimensions of ASFV associated lesions in the field are only very sparsely represented urging more thorough investigations.

Based on this, we aimed to perform pathological examination of wild boar carcasses infected with ASFV to gain more profound knowledge of the pathology of animals succumbing to ASF under natural conditions. We took the opportunity to analyze whether three different variations of the emerging virus variants in Germany may have an impact on the virulence of ASFV and the severity and duration of the disease. For this purpose, detailed pathological and molecular virological investigations were performed on wild boar carcasses infected with variants II, III and IV found in LOS, MOL and SN, respectively.

## Material and methods

### 1. Study design

In accordance with the Animal Disease Crisis Unit of the federal states of Brandenburg and Saxony, sixteen wild boar carcasses were obtained from different outbreak areas (n=7 from Landkreis Oder-Spree (LOS), n=5 from Märkisch-Oderland (MOL), n=4 from Spree-Neiße (SN)) between February and March 2021 where ASF virus variants II, III and IV have emerged as published previously (7). The cadavers were transported to the Friedrich-Loeffler-Institut in compliance with national animal disease and hygiene regulations. Wild boar cadavers were examined in pathological and virological detail. Details on the cadaver material including location of origin, detection of virus variant, age, sex, weight, and preservation status are given in Table 1.

**Table 1.** Summary presentation of examined wild boar carcasses from three different German outbreak areas LOS, MOL and SN.

| No | origin | virus variant | age (year) | sex | weight (kg) | preservation | found dead/shot | Anomalies/comments |
| --- | --- | --- | --- | --- | --- | --- | --- | --- |
| 1 | LOS | II | <1 | female | 10 | good | dead | Brachygnathia superior |
| 2 | LOS | II | <1 | female | 30 | good | dead | / |
| 3 | LOS | II | >2 | female | 62 | autolysis advanced | dead | / |
| 4 | LOS | II | <1 | female | 40 | autolysis moderate | dead | / |
| 5 | LOS | II | <1 | female | 31 | good | dead | / |
| 6 | LOS | II | <1 | male | 37 | autolysis moderate | dead | / |
| 7 | LOS | II | <1 | female | 27 | good | dead | / |
| 8 | MOL | III | <1 | female | 22 | good | dead | / |
| 9 | MOL | III | <1 | female | 28 | good | dead | / |
| 10 | MOL | III | <1 | female | 36 | good | dead | / |
| 11 | MOL | III | <1 | female | 38 | autolysis moderate | dead | / |
| 12 | MOL | III | <1 | female | 36 | autolysis moderate | shot | Lung not available |
| 13 | SN | IV | <1 | male | 36 | good | dead | / |
| 14 | SN | IV | <1 | male | 30 | autolysis moderate | dead | Scavenger feeding marks (thorax) |
| 15 | SN | IV | <1 | female | 31 | good | dead | / |
| 16 | SN | IV | >2 | female | 75 | autolysis moderate | dead | / |

#### 1.1 Pathological examination

##### Necropsy

Full necropsy was performed on wild boar cadavers (n=16). Organ lesions were scored from 0 to 3 (0= normal, 1= mild, 2= moderate, 3= severe; unless not otherwise stated) as recently published (25) with additional modifications shown in Table 2. Tissues samples including popliteal lymph node, spleen, lung, kidney, liver, heart, brain (cerebellum and cerebrum) and adrenal gland were taken from wild boar and fixed in 10% neutral-buffered formalin for at least 3 weeks.

**Table 2.**
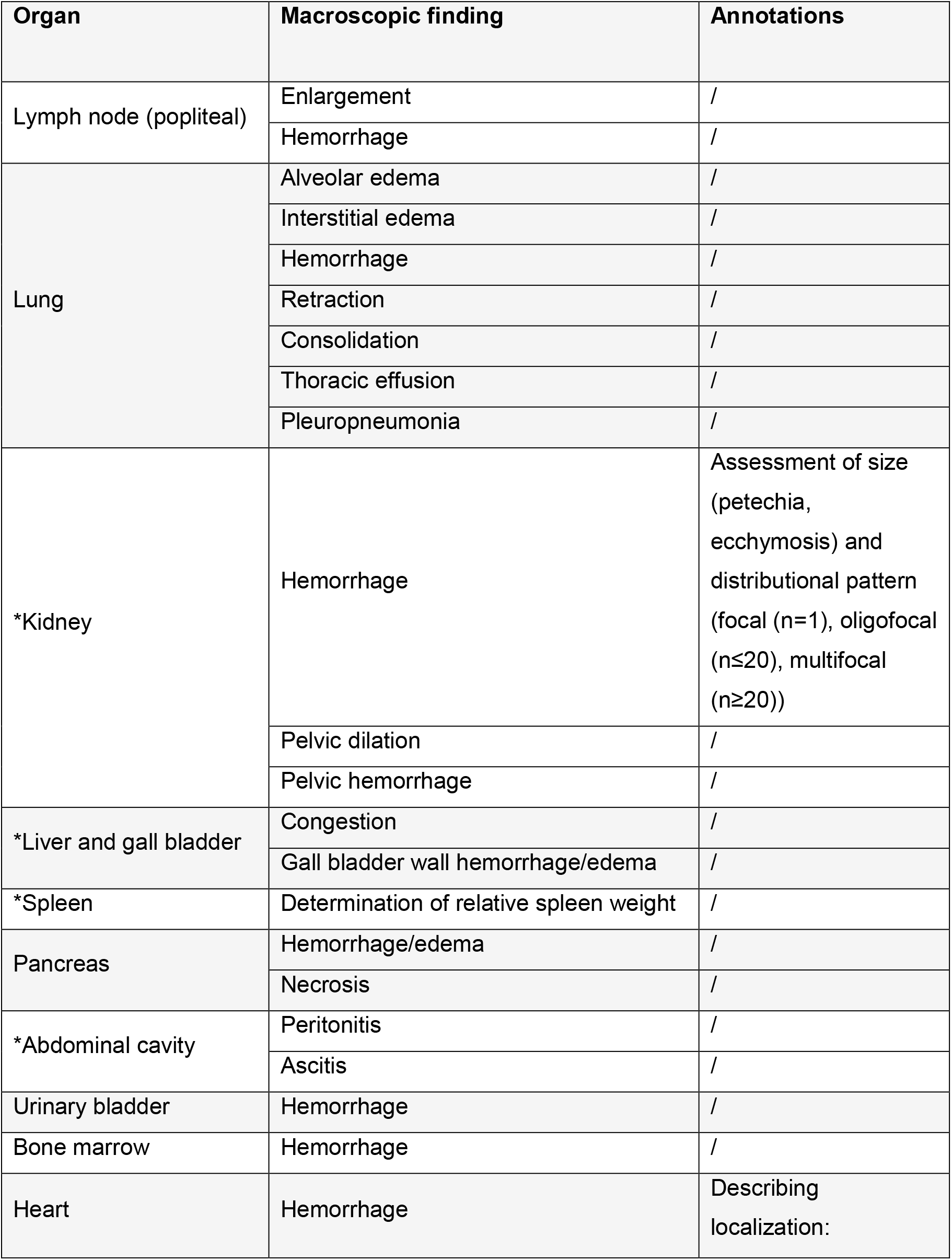
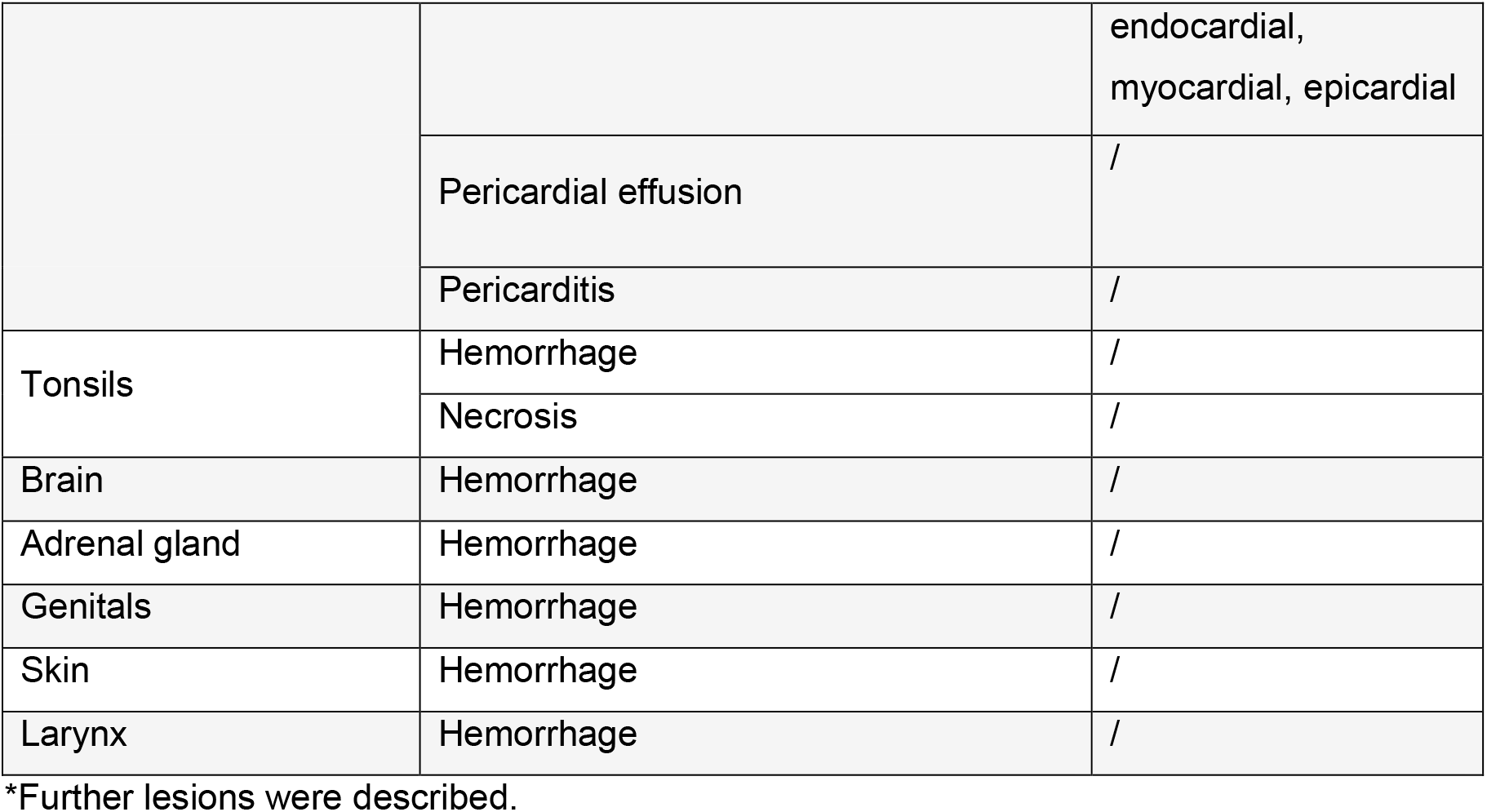
Assessment of gross pathological criteria in ASFV infected wild boar.

##### Histopathology and immunohistochemistry

Tissue samples were embedded in paraffin wax and cut at 2-3 μm slices. Hematoxylin-eosin (HE) staining was performed to examine the main macroscopic lesions in more histological detail. To visualize viral antigen, anti-ASFV p72 immunohistochemistry was conducted on the respective organs as described earlier (17, 18). In brief, sections were treated with an in-house rabbit polyclonal primary antibody against the major capsid protein p72 of ASFV (diluted in TBS 1:1600, 1 h), followed by incubation with a secondary, biotinylated goat anti-rabbit IgG (Vector Laboratories, Burlingame, CA; diluted in TBS in 1:200, 30 min). Positive antigen detection was visualized by the Avidin-Biotin Complex (ABC) method providing horseradish peroxidase that converted the added chromogen 3-amino-9-ethylcarbazole (AEC) into insoluble red-colored deposits at the reaction site. As negative control consecutive sections were labelled with an irrelevant antibody (M protein of Influenza A virus, ATCC clone Hb64). An ASF positive control slide was included in each run.

##### Histopathology including semiquantitative antigen scoring

Slides were scanned using Hamamatsu S60 scanner and evaluated using NDPview.2 plus software (Version 2.8.24, Hamamatsu Photonics, K.K. Japan). While histopathological lesions obtained on HE-stained sections were described only qualitatively (present/absent) due to autolysis-related limited assessability, the viral antigen content in the respective organ was determined on a semiquantitative scoring scale as previously published (18). The most affected area (420×260μm) per sample sections was scored with score 0 (no antigen), score 1 (1–3 positive cells), score 2 (4–15 cells), and score 3 (>16 cells). Cells with fine granular cytoplasmic labelling were considered positive whereas chromogen aggregations without cellular association were not counted.

#### 1.2 Detection of ASFV genome

To determine the viral genome load, tissue samples were homogenized in 1 ml of phosphate buffered saline with a metal bead using a TissueLyzer II (Qiagen GmbH). Viral nucleic acids were extracted from blood and homogenized spleen, lung, liver, kidney, popliteal lymph node, and brain with the NucleoMag Vet Kit (Machery-Nagel) on the KingFisher extraction platform (Thermo Scientific). Quantitative real-time PCR (qPCR) was conducted according to the protocol published by King, Reid (26) with an in-house full virus standard for determination of genome loads on a C1000 thermal cycler with the CFX96 Real-Time System (Biorad).

#### 1.3. Detection of anti-ASFV antibodies

For investigation of ASFV-specific antibodies, an accredited indirect immunoperoxidase test (IPT) was applied according to the standard protocol SOP/CISA/ASF/IPT/1 provided by the European Reference laboratory for ASF with modifications regarding cell and virus type (https://asf-referencelab.info/asf/images/ficherosasf/PROTOCOLOS-EN/2021_UPDATE/SOP-ASF-IPT-1_2021.pdf). As sample material, plasma was obtained from EDTA blood by centrifugation at 18.000 rcf for 10 minutes from German wild boar cadavers and domestic pigs infected with ASFV “Estonia 2014” from a previous trial for comparison. Titers were determined semi-quantitatively by end point dilution from 1:40 to 1:12800.

#### 1.4 Statistical analysis

Using GraphPad Prism (Version 8.4.2) statistical analysis was conducted to determine overall group differences in terms of viral genome load, viral antigen amount, macroscopic lesion scores and antibody titers. For this purpose, the non-parametric Kruskal-Wallis test with post-hoc Dunn’s test was performed. A *p* value ≤0.05 was considered significant.

## Results

### Pathogen detection in blood and tissues

Full necropsy was performed on all wild boar obtained from the outbreak areas LOS, MOL and SN. To determine the amount of viral genome and antigen, spleen, lung, liver, kidney, popliteal lymph node, brain and blood were collected and prepared for qPCR and anti-ASFV-p72 immunohistochemistry, respectively. Positive detection of viral antigen was evaluated based on a semi-quantitative scale as previously applied (18). Details are given in Supplementary Table 1 and 2 (Table S1 and S2) and results are shown in Fig 1.

**Fig 1.**
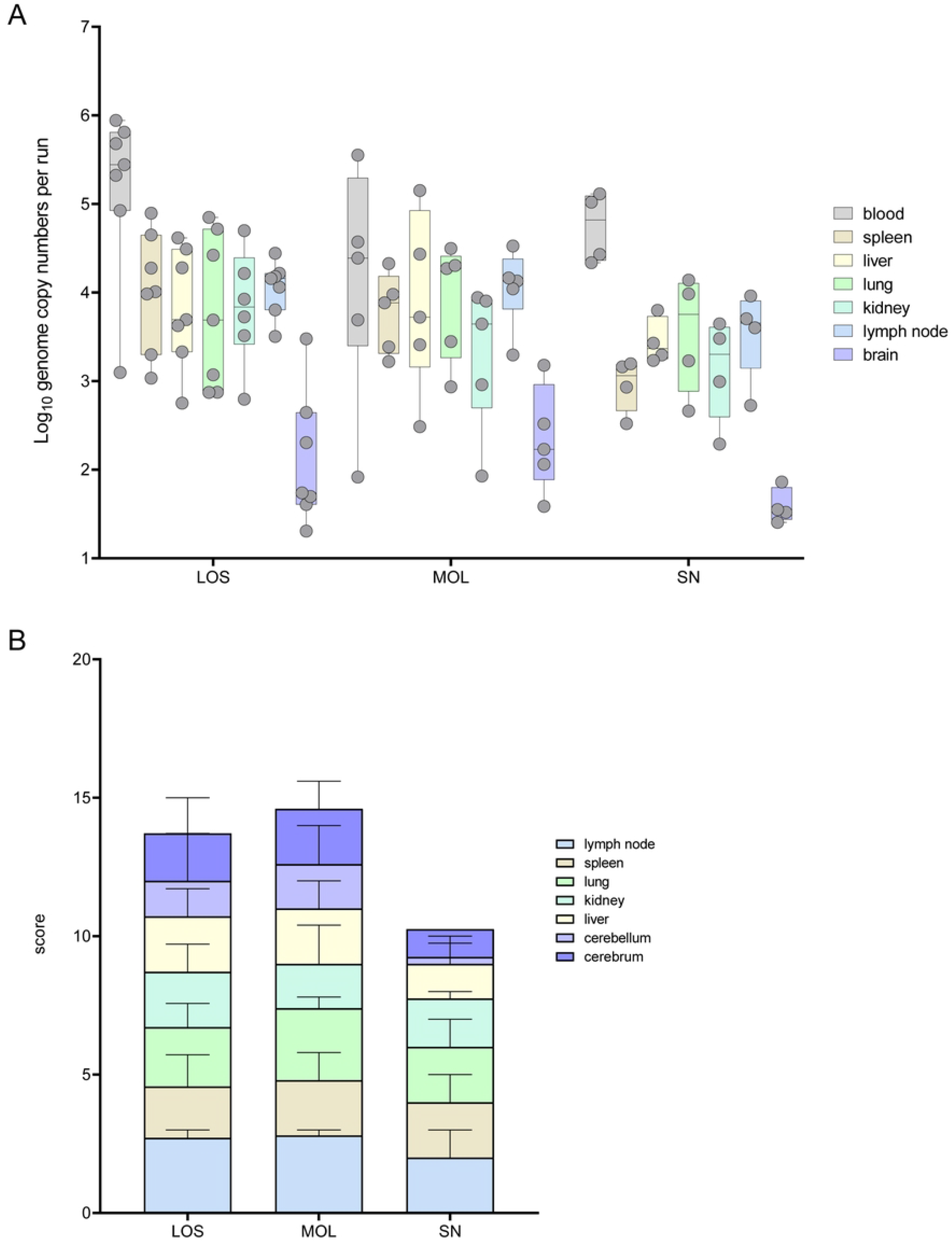
Pathogen detection in blood and tissue samples of ASFV infected wild boar cadavers from LOS, MOL and SN. A) Box plot presenting the individual viral genome load in blood and organ samples. B) Corresponding stacked bar diagram showing the median viral antigen score with range per organ. Organs were scored on a scale from 0 to 3 based on the number of positively labeled cells in the most affected tissue area per high power field.

Viral genome could be found in all samples of infected wild boar. In general, highest viral genome loads were detected in blood samples, varying between 1×10^2^ and 9×10^5^ genome copies (gc) / 5 μl nucleic acid. In general, genome loads in most organ samples were roughly one logarithmic step lower than the corresponding blood samples. In descending order and irrespective of the variant, the viral genome load decreased from spleen, to the lung, kidney, popliteal lymph node, and brain samples. A lower mean viral genome load was detected in wild boar found in SN when compared to animals from LOS and MOL (Fig 1A).

The viral antigen score of selected tissue sections reflected the results obtained by qPCR. Consistent with the lower number of viral genome copies, wild boar from SN reached also lower viral antigen scores (Fig 1B). Details on immunohistochemistry are included in the histopathological evaluation of organ systems in the following section.

### Pathological assessment of organ systems

During necropsy the carcasses were scored macroscopically based on a standardized scoring system (25) with further modifications as indicated in Table 2. In order to investigate macroscopic lesions in more detail, FFPE tissue sections of selected organs (popliteal lymph node, lung, kidney, liver, spleen, heart, brain, adrenal gland) were stained with hematoxylin-eosin. Histopathological alterations were reported only as present/absent due to the reduced number of well-preserved available tissues.

The overall score obtained upon macroscopical evaluation turned out to be the opposite when compared to the viral genome load and antigen score. Therefore, wild boar from SN tended to show a higher total score when compared to wild boar from LOS and MOL (Fig 2). The total score of the animals from MOL was intermediate between the other two groups. A summary of macroscopical findings with individual animal scores are given in Supplementary Table 3 (Table S3).

**Fig 2.**
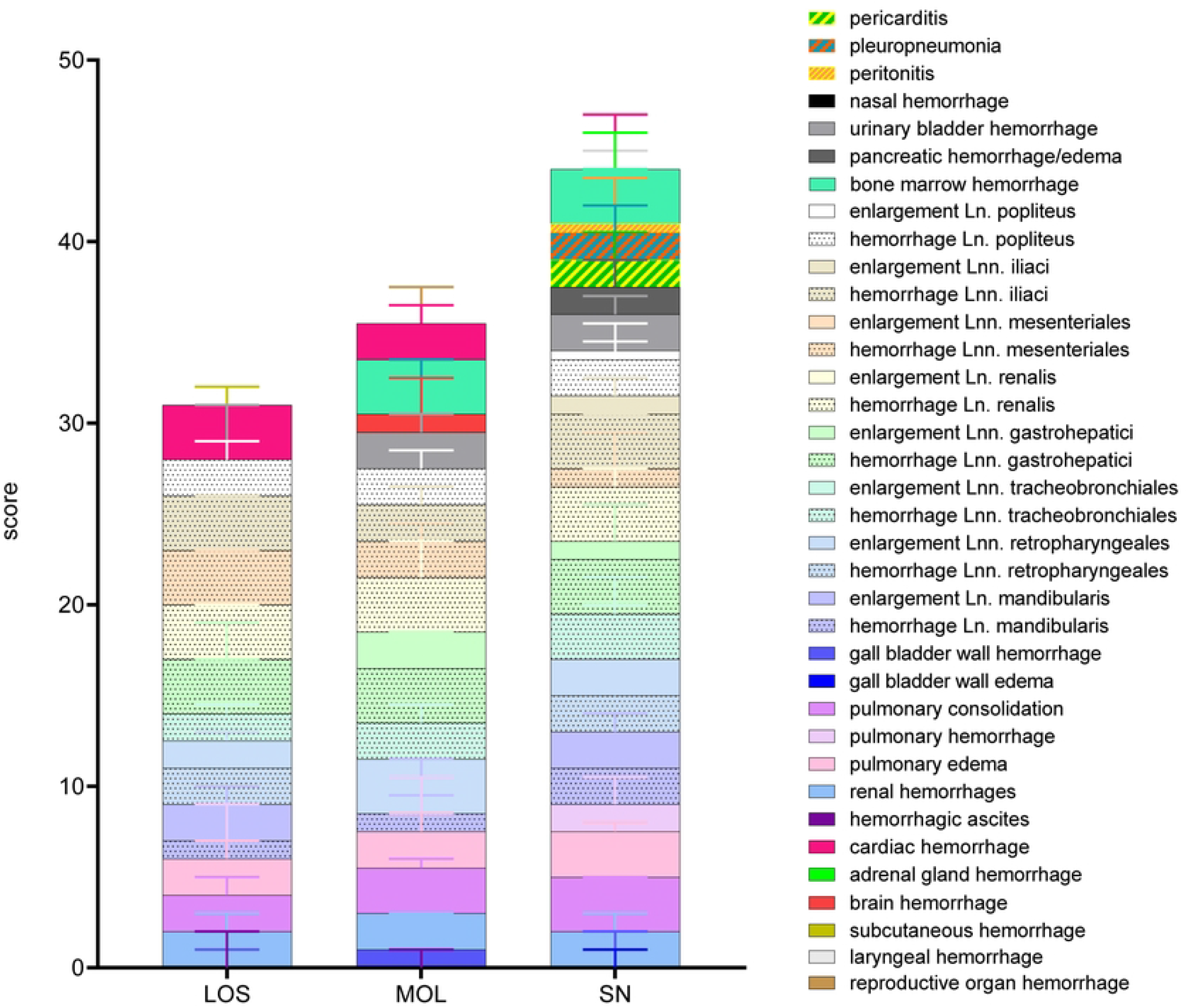
Summary of scoring results following macroscopical investigation of ASFV infected wild boar cadavers from LOS, MOL and SN. Stacked bar diagram showing the total gross lesion score which is composed of individual scores given for macroscopical findings shown on the right. Lesions were scored on a scale from 0 to 3. Bars indicate the median with range.

In the following, detailed gross and histopathological findings of the different organ systems of wild boar found in LOS, MOL and SN will be described.

### Immune system

#### Lymph nodes

##### Gross pathology

For lymph nodes including the mandibular, retropharyngeal, tracheobronchial, gastrohepatic, renal, mesenteric, iliac, inguinal and popliteal nodes details are given in Table S3. An overview of macroscopic scores given for respective lymph nodes as well as representative pathological alterations are shown in Fig 3.

**Fig 3.**
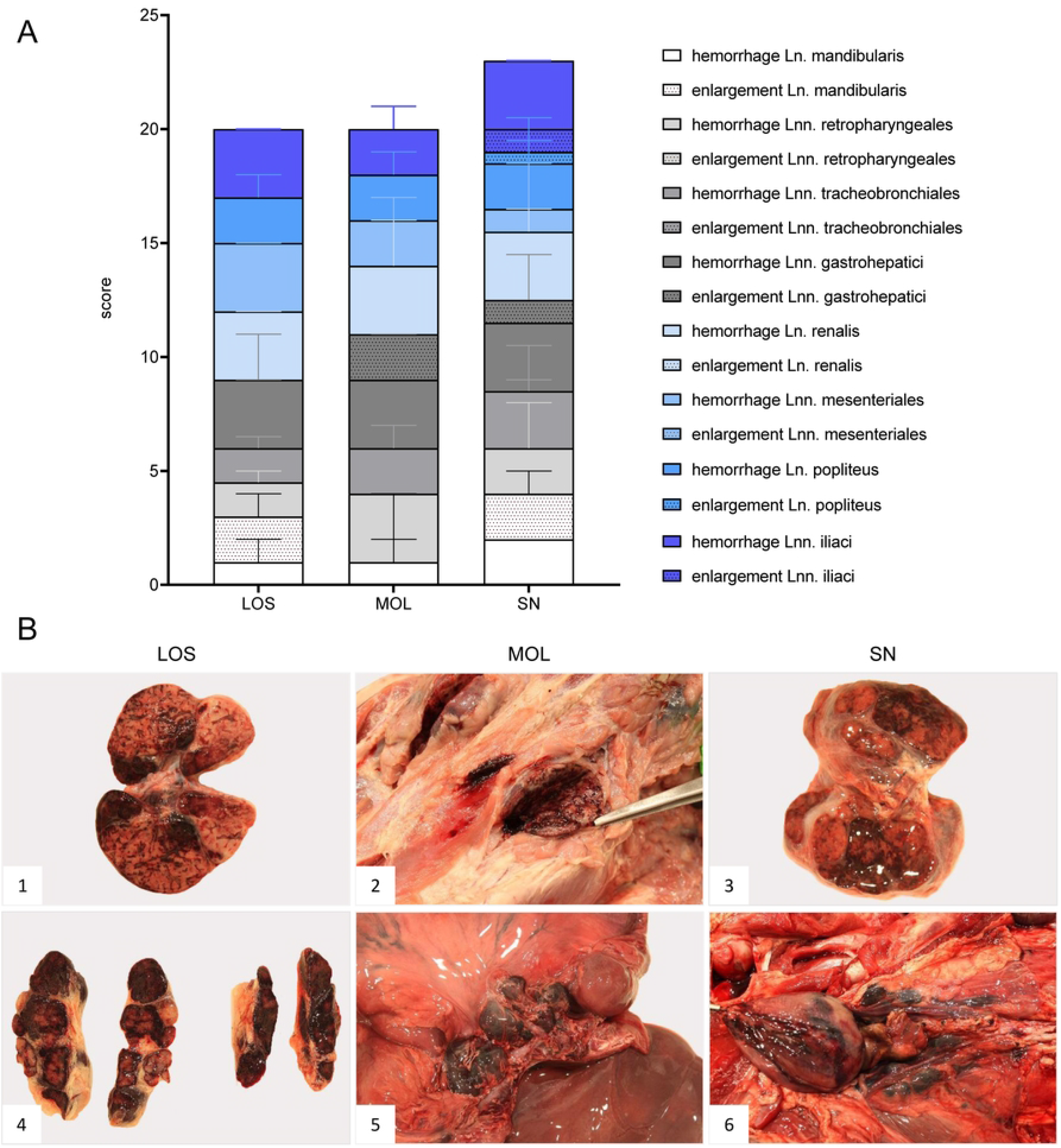
Representative macroscopical findings of lymph nodes in ASFV infected wild boar carcasses from German outbreak areas LOS, MOL and SN. A) Stacked bar diagram showing the total gross lesion score given for enlargement and hemorrhages of various lymph nodes evaluated on a scale from 0 to 3. Bars indicate the median with range per finding. B) Lymph nodes (Ln. mandibularis (1–3), Ln. renalis (4), Lnn. gastrohepatici (5), Lnn. iliaci (6) revealed hemorrhages of varying degree.

In general, wild boar originating from the LOS district showed mild to severe enlargement and hemorrhages, especially of the gastrohepatic and renal lymph nodes followed by mesenteric and iliac nodes. Similar, but generally moderate to severe lesions mainly affecting the gastrohepatic, renal and iliac nodes were observed in wild boar from the MOL and SN area.

##### Histopathology

Since peripheral lymph nodes were best preserved, the popliteal lymph node was examined in more histological detail as demonstrated in Fig 4 and Table S3.

**Fig 4.**
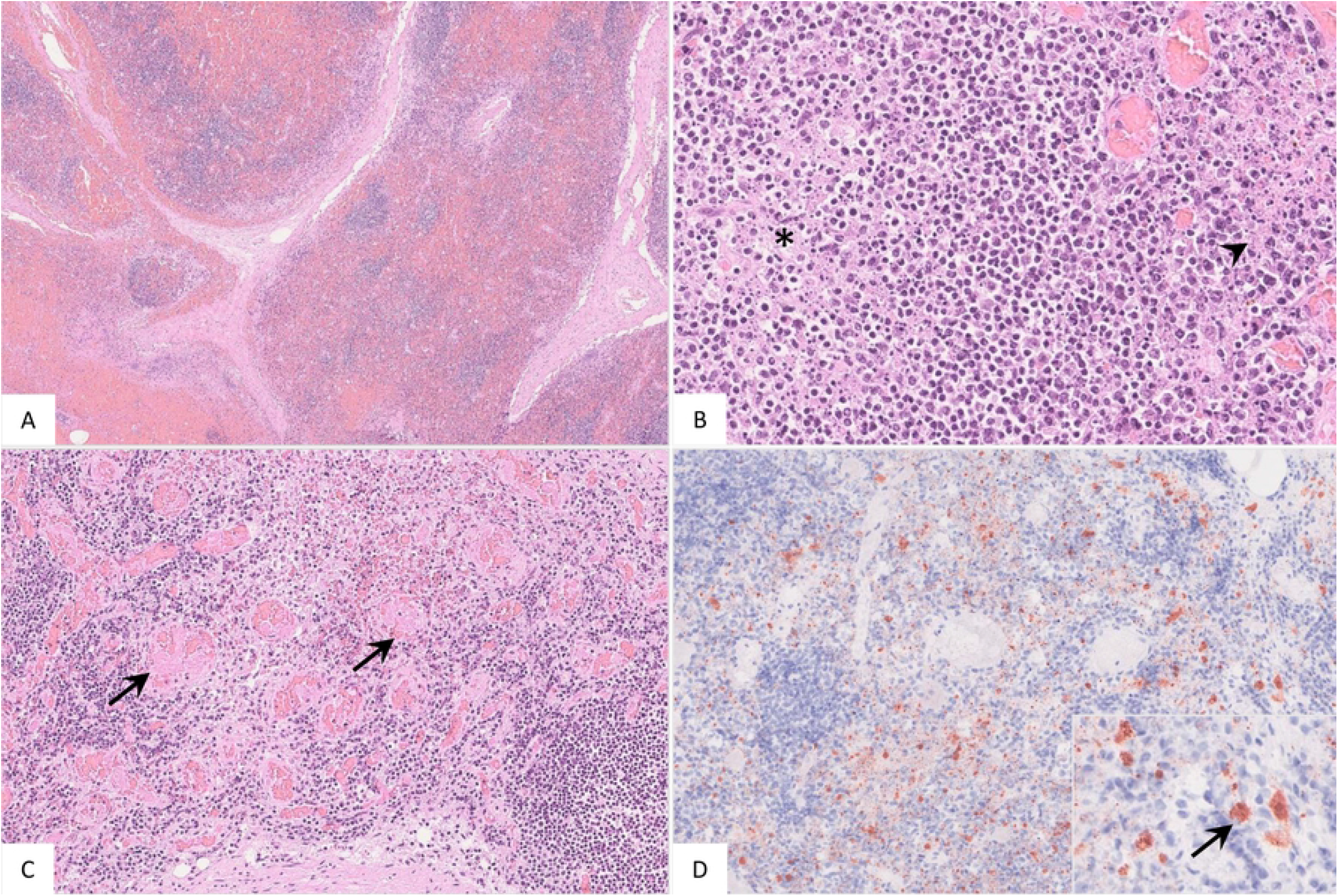
Pathohistological findings of the popliteal lymph node in German ASFV infected wild boar. A) Diffuse lymphoid depletion and hemorrhage affected the follicles, paracortex and medullary chords thereby effacing the physiological lymph node architecture, HE stain. B) A lymphoid follicle with lymphoid depletion (asterisk) was surrounded by necrosis (arrowhead), HE. C) Numerous vessels were occluded by fibrin thrombi (arrows) throughout the lymph node, HE. D) A large number of viral antigen positive cells is shown which were morphologically consistent with macrophages (inlay), anti-p72 immunohistochemistry, ABC method.

Irrespective of the outbreak district lymphoid depletion was found in the lymphoid follicles as well as in the interfollicular areas of the popliteal node. Likewise, diffuse hemorrhages affected cortical follicular, interfollicular and paracortical zones and the medullary cords (Fig 4A). Necrotic areas were observed in the interfollicular areas, paracortex and medullary chords (Fig 4B). Thrombosed vessels were mainly observed in wild boar found in LOS (Fig 4C). Except for one wild boar from SN, all animals showed moderate to high numbers of p72 positively labeled cells morphologically consistent with macrophages (Fig 4D).

#### Spleen

##### Gross pathology

Since the spleen of most wild boar carcasses was poorly preserved, macroscopic assessment was limited. Therefore, the spleen was evaluated by determination of the relative spleen weight shown in Fig 5.

**Fig 5.**
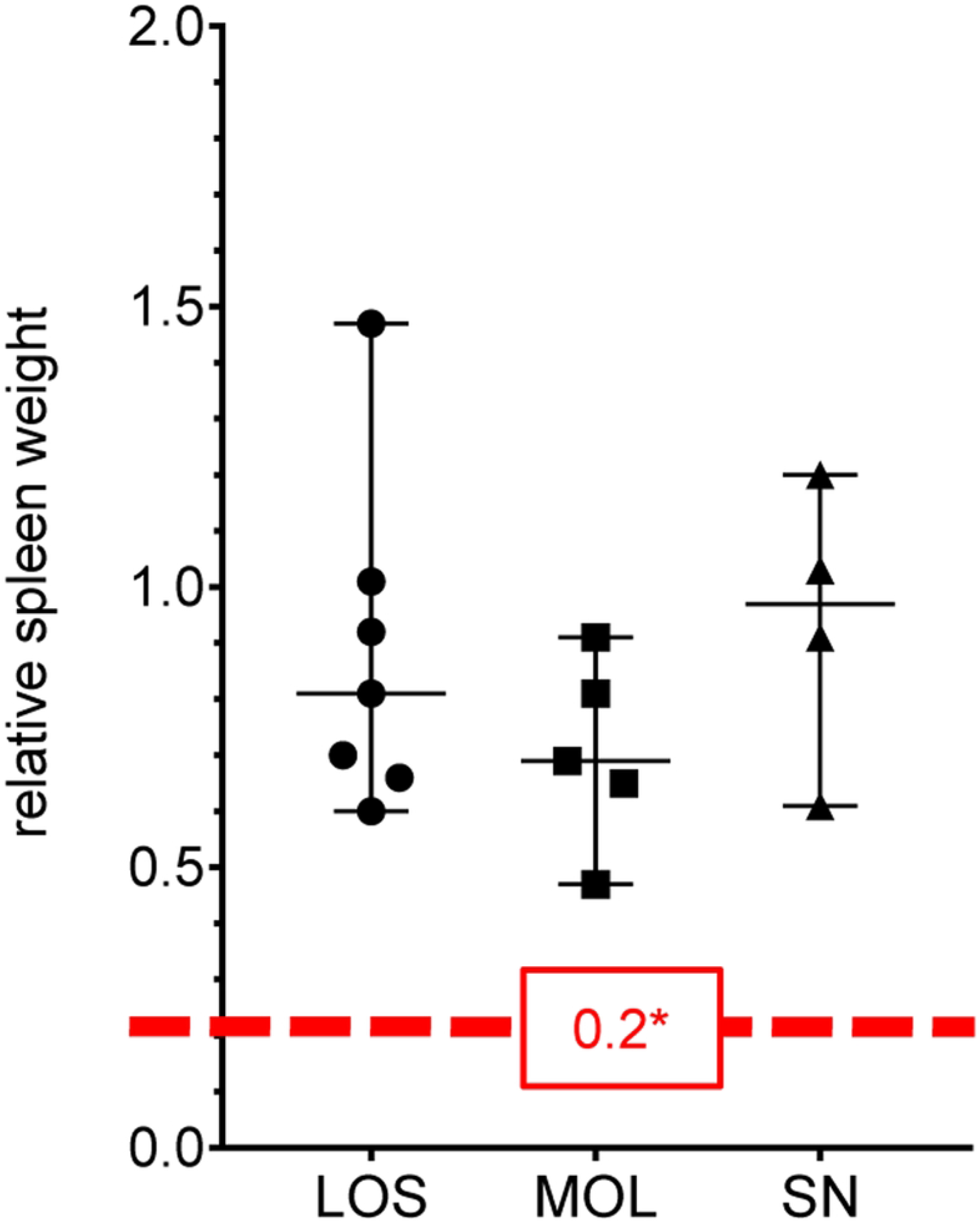
Relative spleen weights of naturally ASFV infected wild boar. Dot plot showing the relative spleen weight values obtained from ASFV infected wild boar from German districts LOS, MOL and SN. The median with range is indicated by error bars. *As reference value the mean relative spleen weight of healthy domestic food-producing pigs of 0.2 was used according to a recent publication (27).

The following findings are based on the physiologic median of relative spleen weight values which were recently determined in domestic pigs (27). High relative spleen weight values were observed in all wild boar irrespective of the district of origin. Median values reached 0.81 (LOS), 0.69 (MOL) and 0.97 (SN) (Figure 5).

##### Histopathology

Histopathological results of the spleen are summarized in Table S3 and Fig 6.

**Fig 6.**
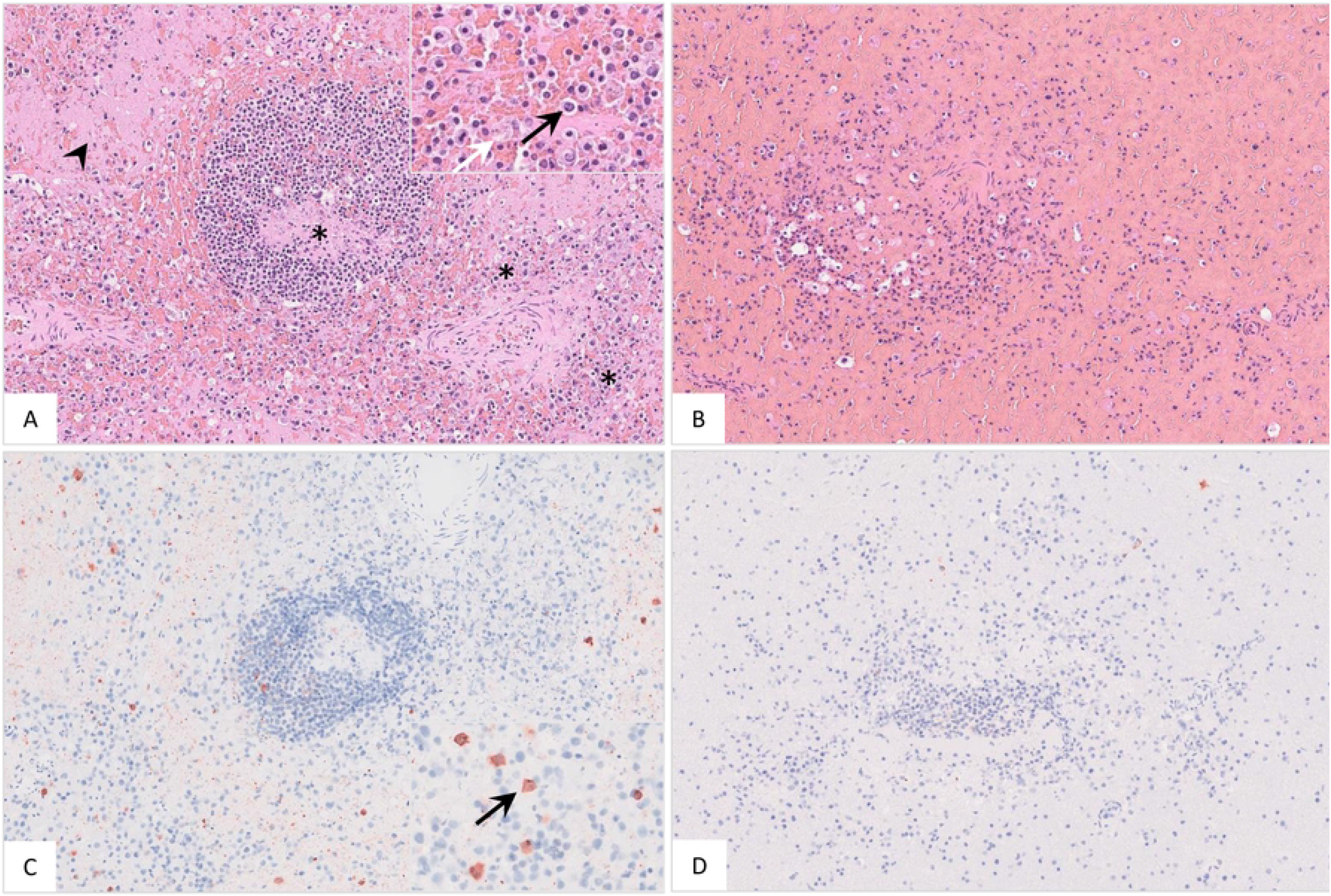
Histopathology of the spleen of naturally ASFV infected wild boar. A) A lymphoid follicle and periarteriolar lymphoid sheaths showed lymphoid depletion (asterisk) with enlarged splenic ellipsoids (arrowhead). Red pulp macrophages were hypertrophic (black arrow) and destructed (white arrow) (inlay), HE. B) Splenic architecture was replaced by necrosis and hemorrhage, HE. C) Mainly moderate, but also D) low numbers of antigen positive cells phenotypically consistent with macrophages (C, inlay, arrow) were present in the spleens, anti p72-immunohistochemistry, ABC method.

Spleen weight determination during macroscopic assessment yielded increased relative spleen weights in all animals regardless of the district in which they were found. Histological examination revealed congestion and hemorrhage in all wild boar. Although the spleen was autolytic in 15 of 16 animals, lymphoid depletion could still be determined. While lymphoid structures including periarteriolar lymphatic sheets (PALS), follicles, and marginal zones were still retained in some animals (Fig 6A), in others the spleen architecture was effaced by necrosis and hemorrhage (Fig 6B). Further, apoptosis/necrosis of myelomonocytic cells of the red pulp was evident in well preserved spleens. The amount of viral antigen highly differed among animals and ranged from low to high numbers of positive cells phenotypically consistent with macrophages (Fig 6C and D). Histopathological results are summarized in Table S3.

#### Bone marrow

##### Gross pathology

Pathological changes of the femoral bone marrow are represented in Fig 7. All scores given for each individual animal can be found in Table S3.

**Fig 7.**
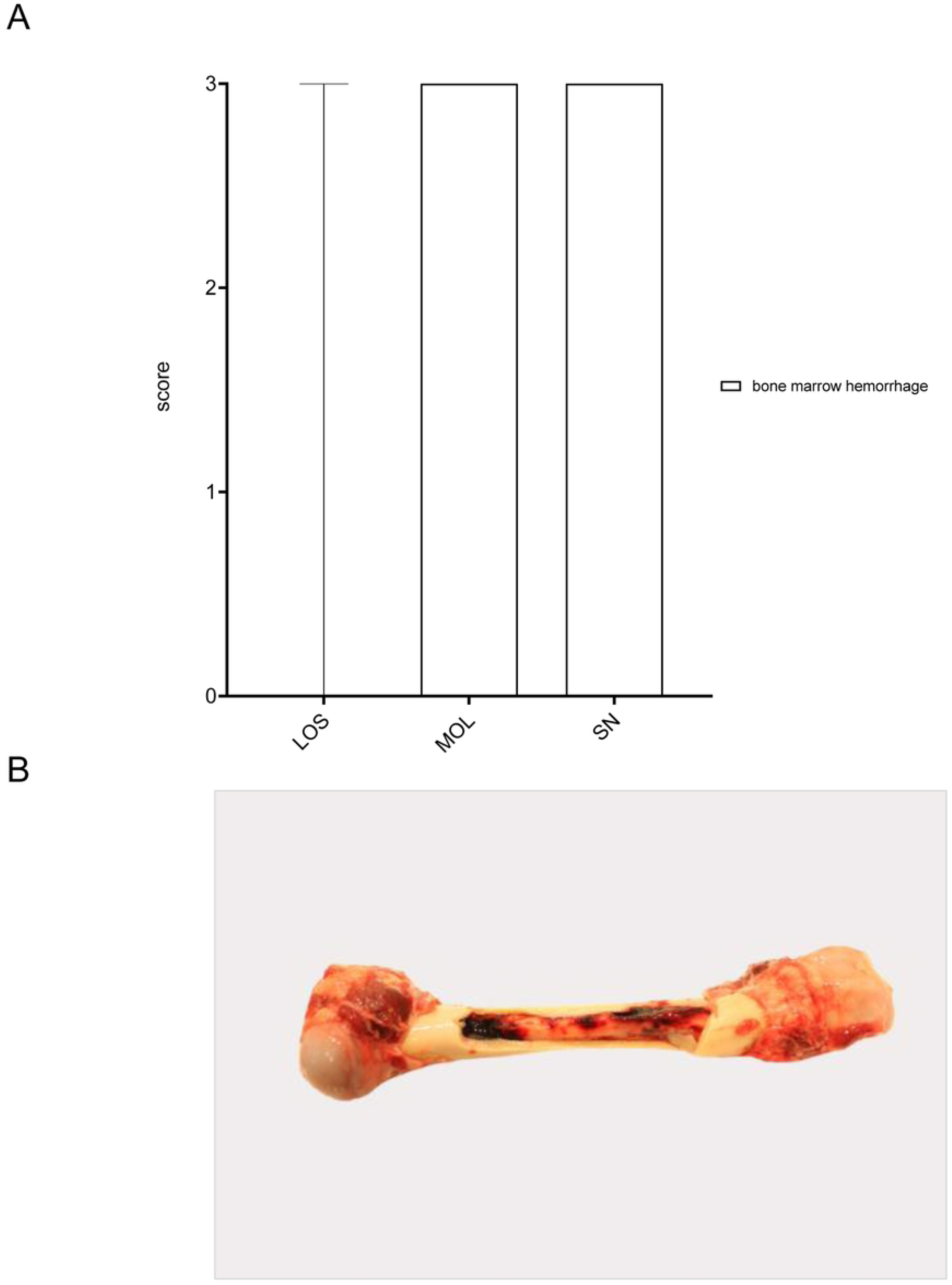
Gross pathology of the bone marrow of naturally ASFV infected wild boar from LOS, MOL and SN. A) Bar diagram showing the scores given for bone marrow hemorrhages on a scale from 0 to 3. Bars represent the median with range. B) Bone marrow hemorrhages, if present, were throughout severe as in this wild boar found in the MOL district.

Three out of seven animals from LOS showed widespread severe hemorrhages in the bone marrow whereas marked lesions were consistently present in all animals from MOL and SN. Histopathological examination was not performed, because in the majority of animals, progression from red to yellow marrow had already occurred in the femur.

### Respiratory system

#### Lung

##### Gross pathology

Details on macroscopic lung scores are summarized in Table S3. Macroscopical scoring and gross findings of lungs of ASFV infected wild boar are represented in Fig 8.

**Fig 8.**
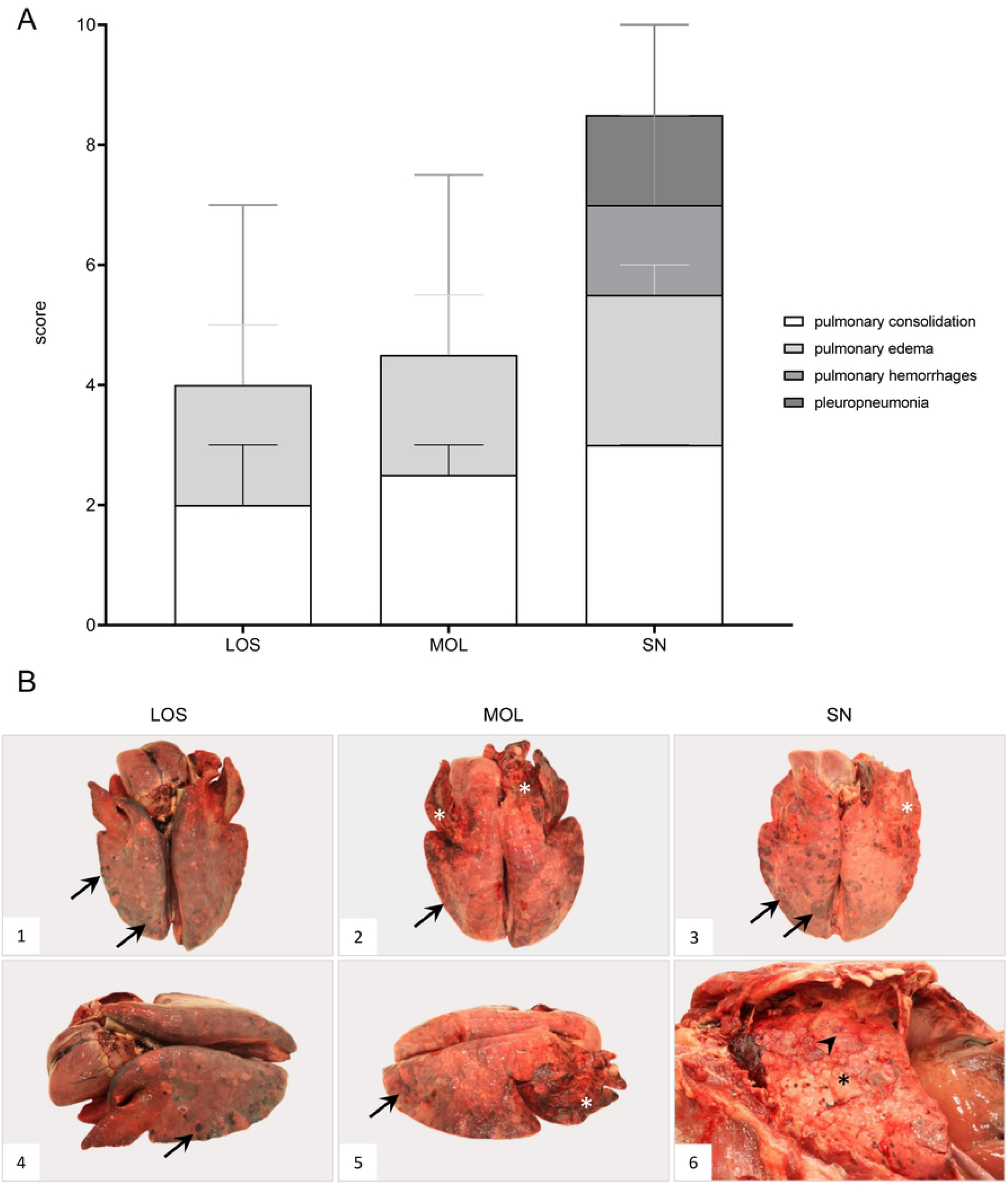
Macroscopical lung lesions of ASFV infected wild boar found in the German districts LOS, MOL and SN. A) Stacked bar diagram demonstrating the median with range of individual scores given for each pathological criterium shown on the right legend. The presence and severity of each finding was scored from 0 to 3. B) All lungs showed consolidated areas of different size (asterisk) and loss of retraction (B1-6). Pulmonary hemorrhages of varying severity are demonstrated by arrows (B1-5). Chronic pleuropneumonia is shown in B5-6 with extensive fibrous pleural adhesions (arrowhead).

Briefly, in wild boar from LOS, moderate to severe pulmonary alveolar edema was present in 5/7 animals. In all wild boar incomplete retraction with consolidation of varying severity was found. Congestion was present in all wild boar while widespread hemorrhages were observed in 3/7 animals. In one animal, severe, focal, fibrinous, chronic pleuropneumonia was evident in the caudal lobe.

Lung lesions could be assessed in 4/5 animals since one boar was killed by a lung shot. In three animals, alveolar edema was detected of moderate to severe degree. The same animals showed loss of retraction whereas variable consolidation was present in all four affected animals. Two wild boar showed severely consolidated areas, one of these animals further had marked, chronic fibrous pleuropneumonia of the right lung. While congestion varied between the animals, severe hemorrhages were found in two wild boar.

Similar lesions occurred in wild boar found dead in SN. Up to severe alveolar edema was present in all animals. The lung of one wild boar was severely autolytic making any further evaluation impossible. In the remaining three pigs, signs of pulmonary inflammation were evident. All lungs lacked retraction and consolidation was severe in two wild boar. All lungs were severely congested, but extensive hemorrhages were found only in two animals. Severe, chronic fibrinous pleuropneumonia of the right lung was present in two animals.

##### Histopathology

During macroscopic assessment, pulmonary consolidation as well as fibrous pleuropneumonias accompanied by hemorrhages and alveolar edema were found, which, with few exceptions, could be confirmed histopathologically as shown in Fig 9 and Table S2. Pulmonary inflammation showed two main manifestations. On the one hand, fibrino-suppurative to necrotizing bronchopneumonias could be detected in few animals from LOS and MOL (Fig 9A-C). In others, the necrotizing inflammatory process was rather limited to the pulmonary interstitium and bronchioles (Fig 9D). In all animals, except for one wild boar from LOS and two animals from SN moderate to high amounts of positively labeled cells consistent with intravascular, -alveolar and interstitial macrophages were detected by immunohistochemistry (Fig 9E and F).

**Fig 9.**
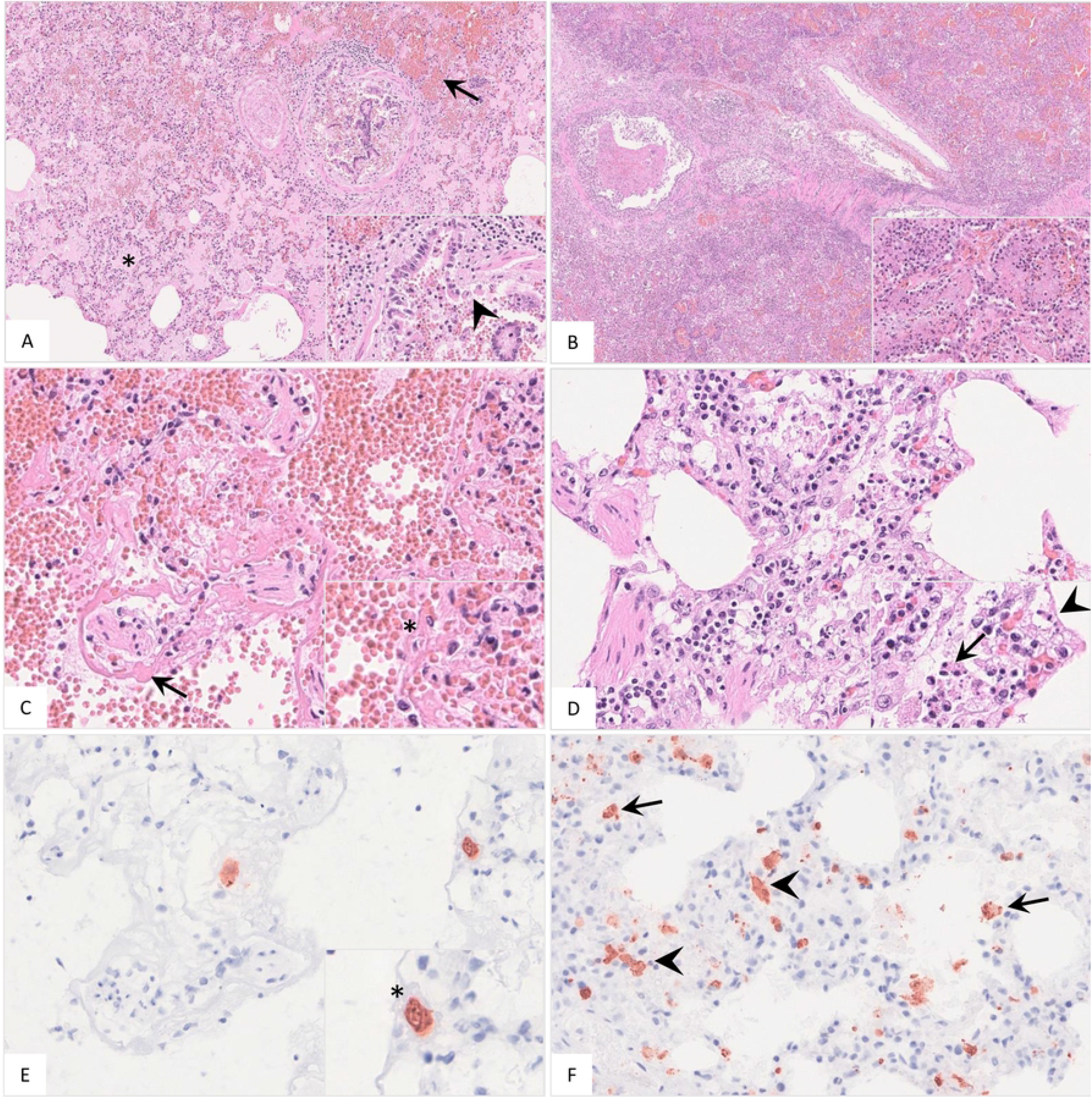
Histopathological findings of lungs in German naturally ASFV infected wild boar. A) Alveoli were filled with protein-rich edema fluid (asterisk), erythrocytes (arrow), and fibrin strands. The bronchiolus revealed epithelial necrosis (inlay, arrowhead) and contained cellular debris and erythrocytes. A distended pulmonary vein with fibrin thrombi was present left from the bronchiole, HE. B) In few animals severe fibrino-suppurative to necrotizing bronchopneumonia was detected. Alveoli were densely filled with cellular debris, fibrin, viable and degenerate neutrophils, plasma cells, macrophages and lymphocytes as well as erythrocytes (inlay), HE. C) A low number of wild boar showed loss of alveolar epithelium and hyaline membranes (arrow). An intravascular macrophage is indicated by asterisk (inlay), HE. D) Interstitial inflammation was noted mainly in animals from SN. Alveolar septa showed epithelial necrosis (inlay, arrowhead), infiltration by necrotic macrophages (inlay, arrow), neutrophils, lymphocytes and plasma cells, HE. E and F) Immunohistochemistry showed viral antigen positive cells morphologically consistent with intravascular (E, asterisk, consecutive section of (C)) intraalveolar (F, arrow) and interstitial (F, arrowhead) macrophages, anti p72-immunohistochemistry, ABC method.

### Cardiovascular system

#### Heart

##### Gross pathology

Macroscopic scoring and representative cardiac lesions are summarized in Fig 10. Individual gross scores are included in Table S3.

**Fig 10.**
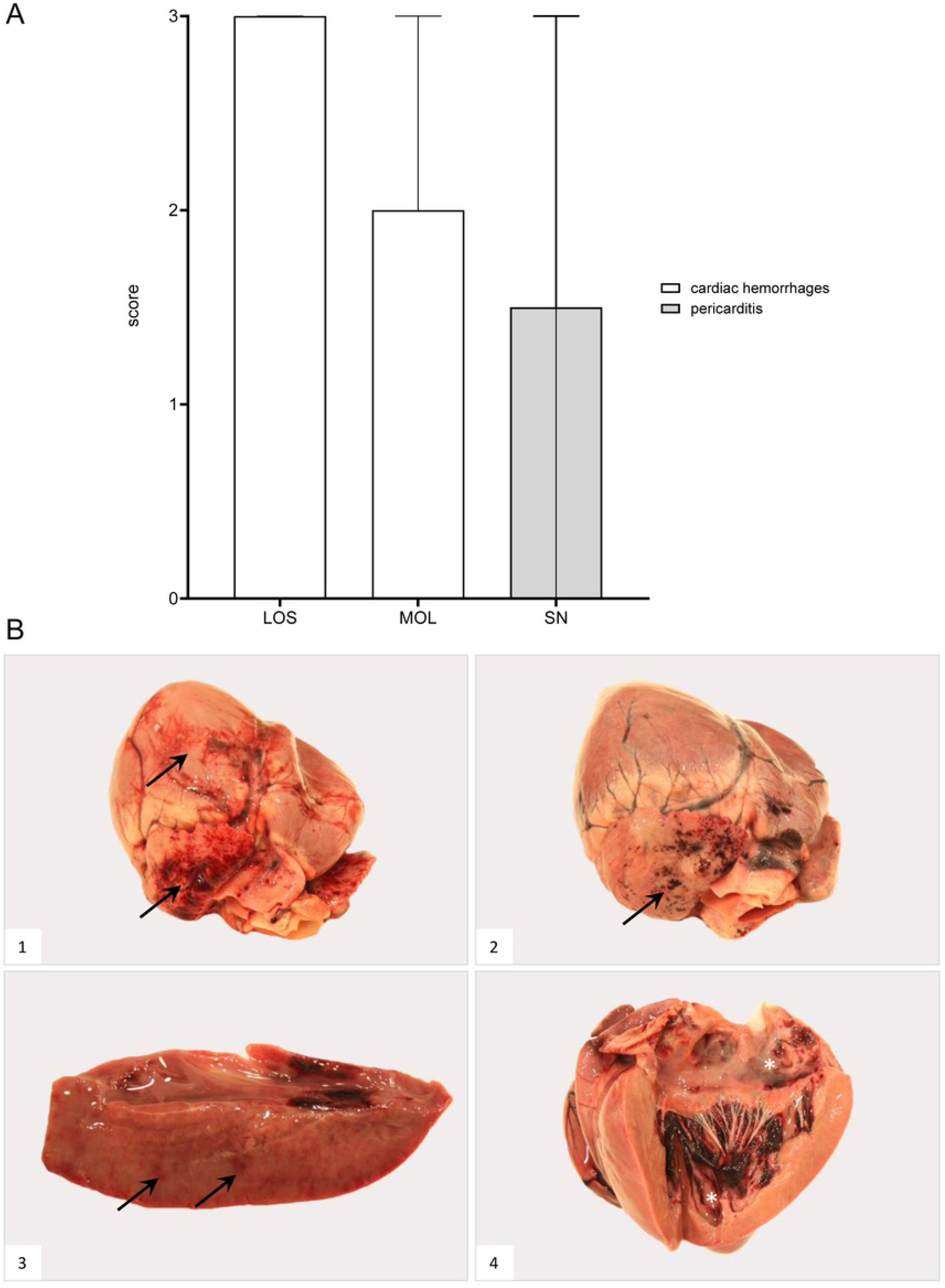
Heart lesions in naturally ASFV infected wild boar from German outbreak areas LOS, MOL and SN. A) Scoring of the heart included the presence and severity of hemorrhages as well as pericarditis which were evaluated on a scale from 0 to 3. Bars indicate the median with range. B) Hemorrhagic lesions of different locations and severity of ASFV infected wild boar found in the LOS district are shown. Multifocal paintbrush to coalescing hemorrhages were found in the epicardium (arrow) to a variable extent (B1-2). Scant myocardial hemorrhages (arrow) are indicated in B3. Multifocal endocardial hemorrhages (asterisk) are present in B4. The darker blood coagulum had to be differentiated from hemorrhages.

Severe hemorrhages were more frequently seen in 5/7 wild boar found in the LOS district. The epicardium was affected in five, the myocardium in three and the endocardium in four animals.

Comparable severe lesions were present in 3/5 wild boar from MOL. Changes were observed in the epi- and endocardium in two animals whereas the myocardium was additionally affected in another wild boar.

In contrast, only 1/4 wild boar from the SN area showed widespread bleeding located to the epi- and myocardium. Extensive, chronic fibrinous pericarditis was further detected in two wild boar from SN.

##### Histopathology

Detailed histopathological evaluation of the heart can be found in Table S2. Representative microscopic lesions are shown in Fig 11.

**Fig 11.**
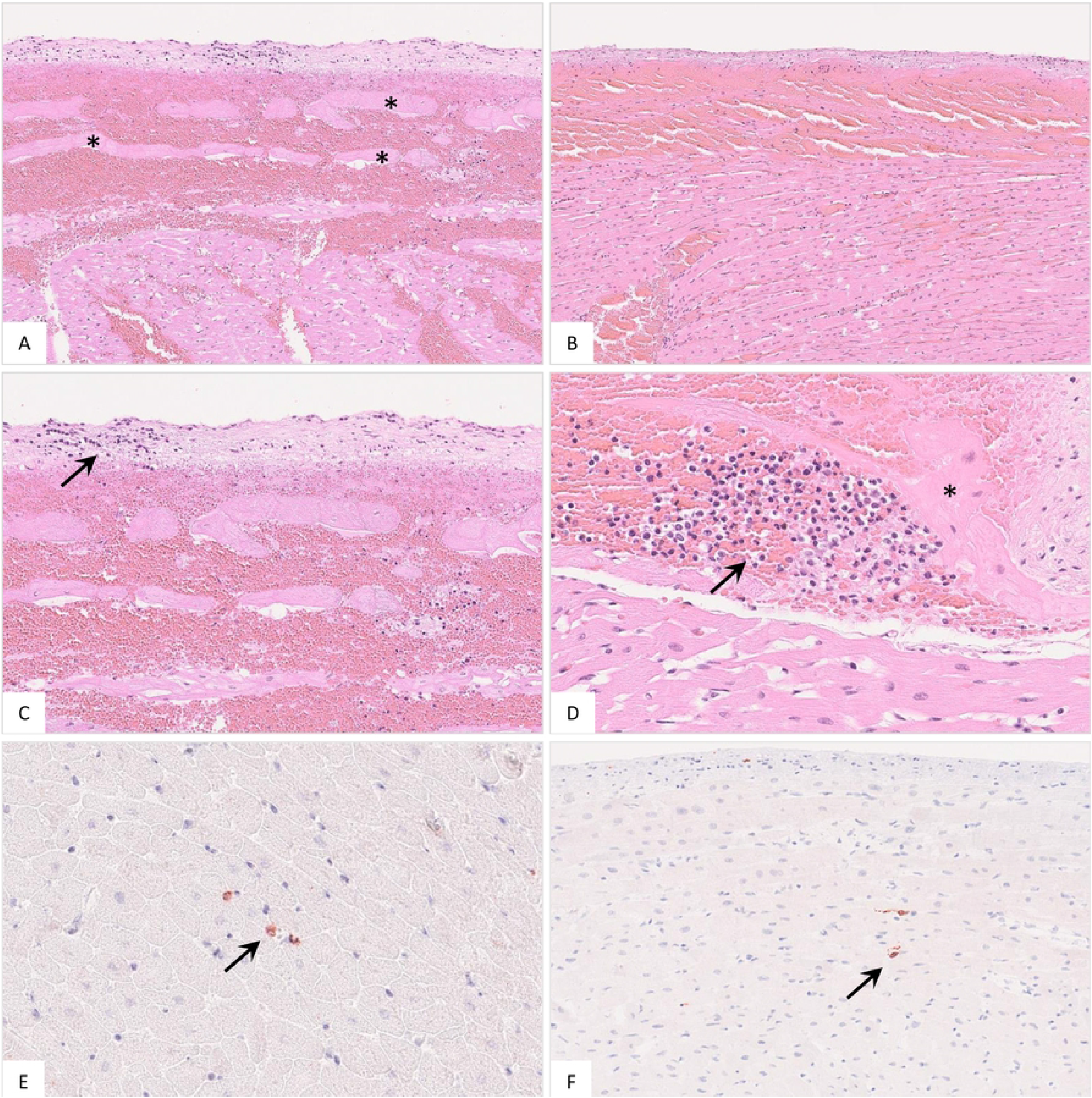
Histopathology of the heart in naturally ASFV infected wild boar from Germany. A) Massive hemorrhage involved the endocardium as well as the myocardium, displacing subendocardial Purkinje fibers (asterisk), HE. B) The epicardium was also affected by diffuse hemorrhage radiating into the myocardium, HE. C) Higher magnification from (A) shows minimal accumulation of infiltrating mononuclear cells in the endocardium (arrow), HE. D) Subendocardial infiltrates (arrow) were also present between Purkinje fibers (asterisk), HE. E and F) Immunohistochemistry of the heart showed only few positive macrophages (arrow), anti-p72 immunohistochemistry, ABC method.

Macroscopic hemorrhages in the epi-, myo-, and endocardium of varying degrees were confirmed histopathologically (Fig 11A and B). In addition, in few animals (n=3), there was endocardial and subendocardial infiltration by mononuclear cells (Fig 11C and D). Viral antigen was found only in a low to moderate number of cells morphologically mostly consistent with macrophages (Fig 11E and F).

### Urinary system

#### Kidney

##### Gross pathology

Detailed scoring of the kidney and gross lesions are shown in Table S3 and Fig 12.

**Fig 12.**
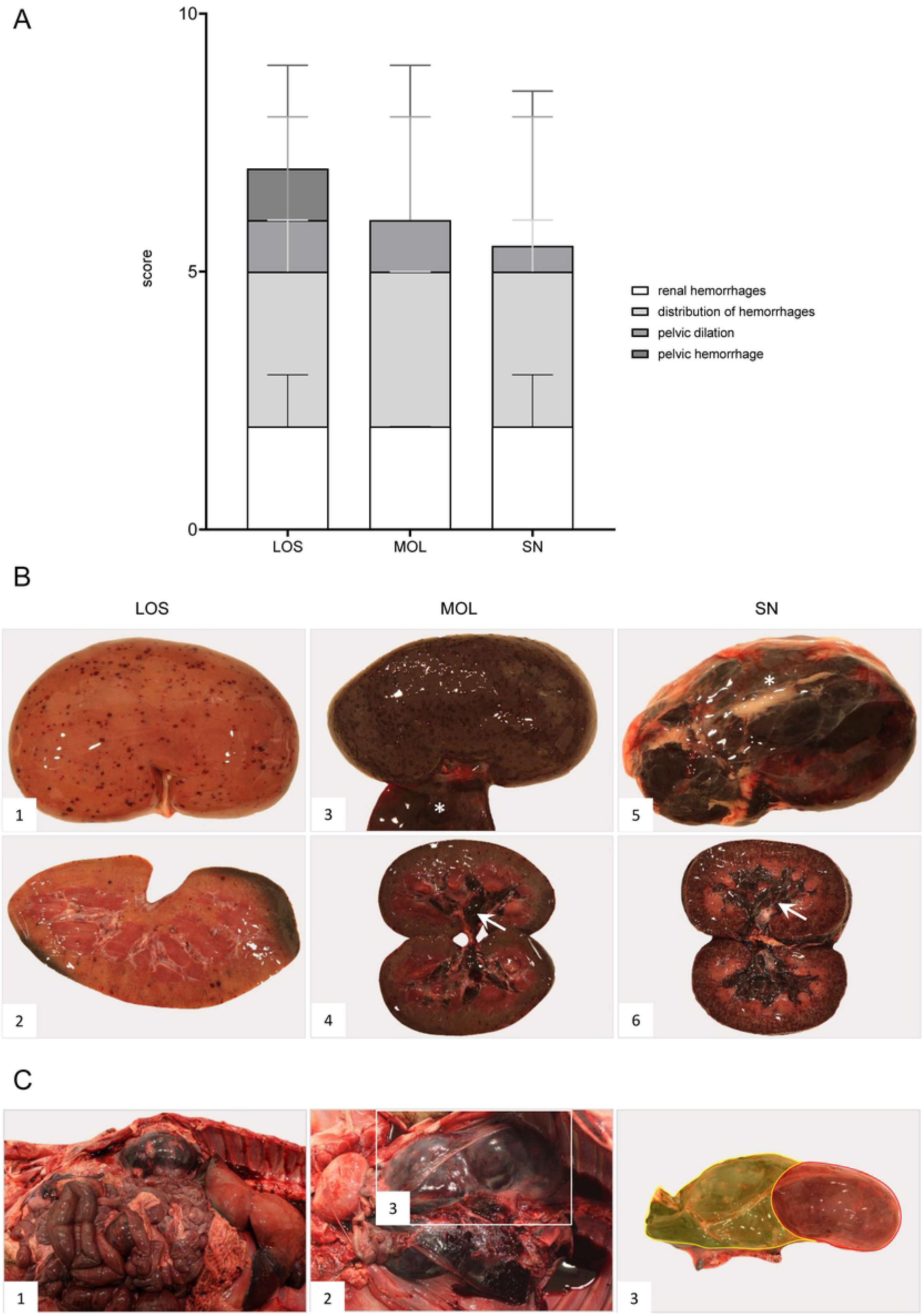
Pathologic changes in kidneys of ASFV infected German wild boar from outbreak areas LOS, MOL and SN. A) Stacked bar diagram of gross lesion scoring of pathological criteria listed on the right. Scoring was conducted on a scale from 0 to 3 or from 0 to 4 (distribution pattern of hemorrhages). Individual scores are given as median values with range. B) Hemorrhagic lesions of various size and severity affecting different parts of the organ are shown in B1-6. Multifocal petechiae with fewer ecchymoses primarily located to the renal cortex are depicted in B1-2. Grey discoloration of the kidney periphery was due to beginning autolysis (B2). Mainly affecting the renal cortex (cortico-medullar pattern), diffuse ecchymoses are present in B3-4. Marked dilation and diffuse bleeding into the renal pelvis are depicted in B4 and B6 (arrows). To a lesser extent, oligofocal pinpoint hemorrhages (arrowhead) could be found in the medulla (B6). Edema of the perirenal tissue is represented in B3 and B5 (asterisk). C) Massive hemorrhage resulted in expansion and bulging of the renal capsule (C1 and C2). The hemorrhage further extended into the perirenal and retroperitoneal tissue including the ureter (C2). To better distinguish the kidney and the extent of hemorrhage from C2, the kidney was shaded red and the hemorrhage was highlighted in yellow (C3).

The kidneys of all wild boar found in LOS were evaluated for hemorrhages. Two wild boar showed focal or multifocal petechiae, three revealed multifocal to diffuse ecchymoses and another two had diffuse hemorrhages throughout the organ. A cortical-medullar pattern was observed in four animals whereas in three pigs hemorrhages were equally distributed in the cortex and medulla. Dilation of the renal pelvis with few hemorrhages was present in four animals. In one animal severe perirenal edema was present.

Renal hemorrhages were detected in 4/5 wild boar from MOL. Three animals showed diffuse ecchymoses, one revealed multifocal petechiae. Except for one animal that presented a cortical-medullar pattern, hemorrhages occurred equally in the cortex and medulla. In two animals there was marked dilation of the renal pelvis with diffuse hemorrhages. Perirenal edema and hemorrhages of marked severity were found in one wild boar.

Renal lesions were observed in 3/4 wild boar from SN. Multifocal to diffuse ecchymoses were detected in two animals whereas in one pig the whole kidney was affected by diffuse hemorrhage. The cortex was more severely affected than the medulla. Mild to severe pelvic dilation in two wild boar with additional diffuse hemorrhages in another animal were seen. In one pig hemorrhages extended to the perirenal tissue massively dilating the renal capsule and periureteral tissue.

##### Histopathology

Details on histopathological findings are shown in Table S2 and representative lesions are depicted in Fig 13.

**Fig 13.**
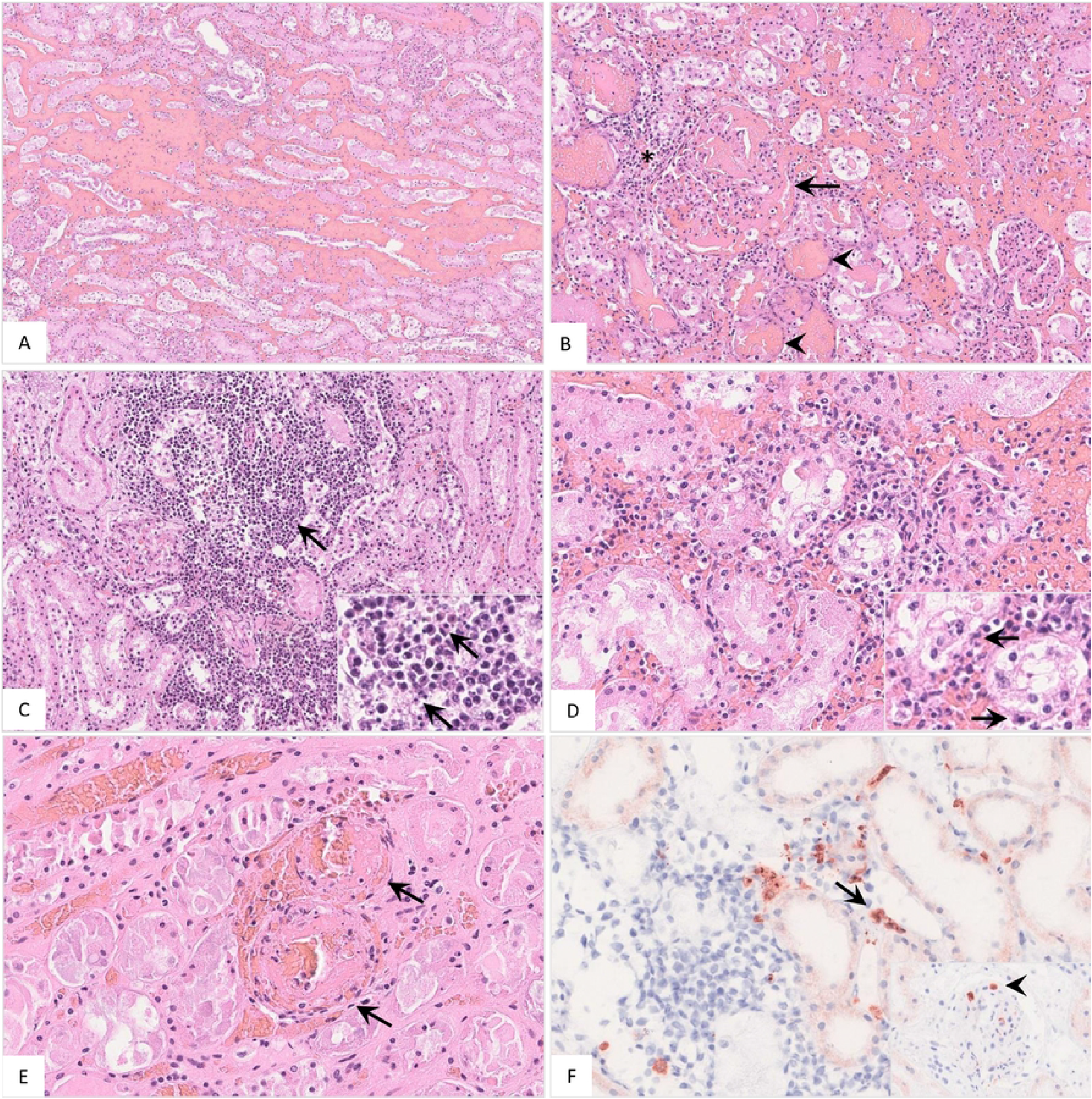
Pathohistological findings of the kidney in naturally ASFV infected wild boar. A) Diffuse hemorrhages were present expanding the renal medullary interstitium, HE. B) A glomerulus showed extravasation of fibrin admixed with erythrocytes into the Bowman’s space (arrow). There was periglomerular infiltration of partly degenerated mononuclear cells (asterisk). Red blood cell casts were present in several tubules surrounding the glomerulus (arrowhead), HE. C) Extensive mononuclear cell infiltrates accumulated around tubules and glomeruli (arrow) and revealed multiple foci of apoptosis/necrosis (inlay, arrow), HE. D) In some areas, tubulointerstitial nephritis was associated with tubular epithelial apoptosis/necrosis (inlay, arrow), HE. E) Fibrinoid vascular necrosis could be found in varying amounts of renal veins (arrow), HE. F) Representative immunohistochemical image showing moderate numbers of positively labeled macrophages in the renal interstitium (arrow) or glomerular capillaries (inlay, arrowhead), anti-p72 immunohistochemistry, ABC method.

Gross examination of the kidneys revealed hemorrhages of varying extent and severity, which histologically were proven to be mainly located interstitially affecting both the cortex and medulla as well as the renal pelvis (Fig 13A). Isolated glomerular lesions with hemorrhages and fibrin deposition in Bowman spaces were also detected (Fig 13B). Except for the animals that revealed progressive autolysis, non-suppurative tubulointerstitial nephritis was diagnosed (Fig 13C). In those animals, tubular epithelial necrosis (Fig 13D) and renal vein thrombi occurred frequently (Fig 13E). Immunohistochemistry revealed varying amounts of positive cells morphologically consistent with macrophages (Fig 13F).

#### Urinary bladder

##### Gross pathology

Gross scoring and representative lesions of the urinary bladder are shown in Fig 14. Individual scores can be found in Table S3.

**Fig 14.**
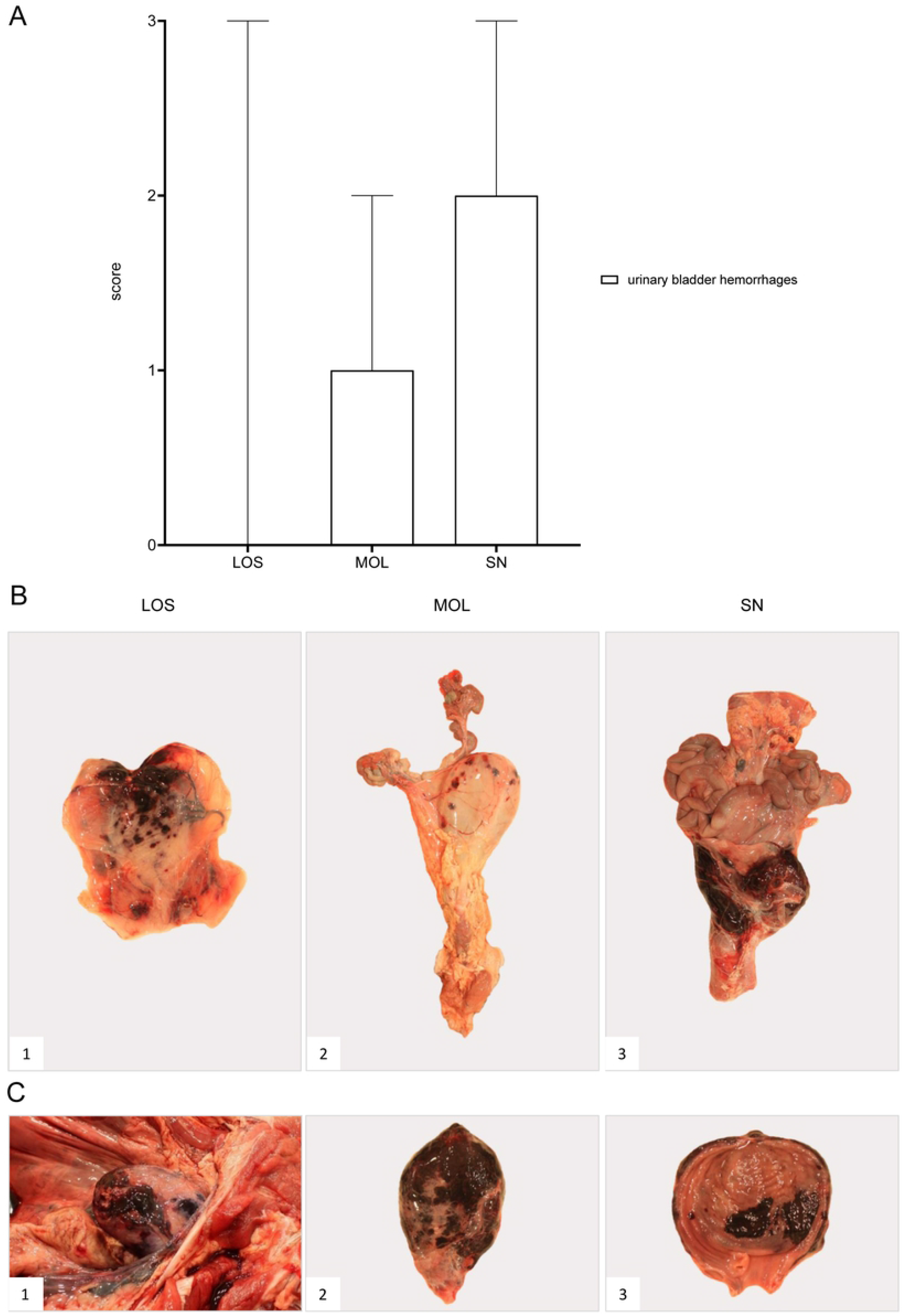
Pathology of the urinary bladder in naturally ASFV infected wild boar from German outbreak districts LOS, MOL and SN. A) Bar diagram showing hemorrhagic changes of the urinary bladder scored on scale from 0 to 3. Bars indicate the median with range. B) Hemorrhages of varying severity were observed during necropsy. Multifocal to coalescing hemorrhages were found only in one pig from LOS (B1). Multiple ecchymoses were present in animals from MOL (B2). In contrast, severe, diffuse hemorrhage of the urinary bladder radiating into surrounding connective tissue was found more frequently in animals from SN (B3). C) Severe hemorrhages were found in wild boar from SN. Extensive, multifocal to coalescing hemorrhages were located to the serosa (C1 and 2) as well as to the mucosal surface of the urinary bladder (C3).

Multifocal to coalescing hemorrhages of the urinary bladder wall were present in one wild boar found in the LOS district while all other animals revealed no lesions.

More frequently, hemorrhagic lesions in the urinary bladder occurred in 3/5 wild boar cadavers from MOL. One wild boar displayed very few mucosal petechia while the two others presented with mottled hemorrhages of moderate severity.

In 3/4 animals originating from the SN district urinary bladder hemorrhages were most severe. Two out of three animals showed large coalescing areas of hemorrhages affecting the serosa and the mucosa. Milder lesions were observed in the other animal.

Histopathological examination was not performed due to poor preservation of the tissue.

### Gastrointestinal system

#### Liver and gall bladder

##### Gross pathology

Macroscopic scoring results of liver lesions and respective images are shown in Fig 15. Details are given in Table S2.

**Fig 15.**
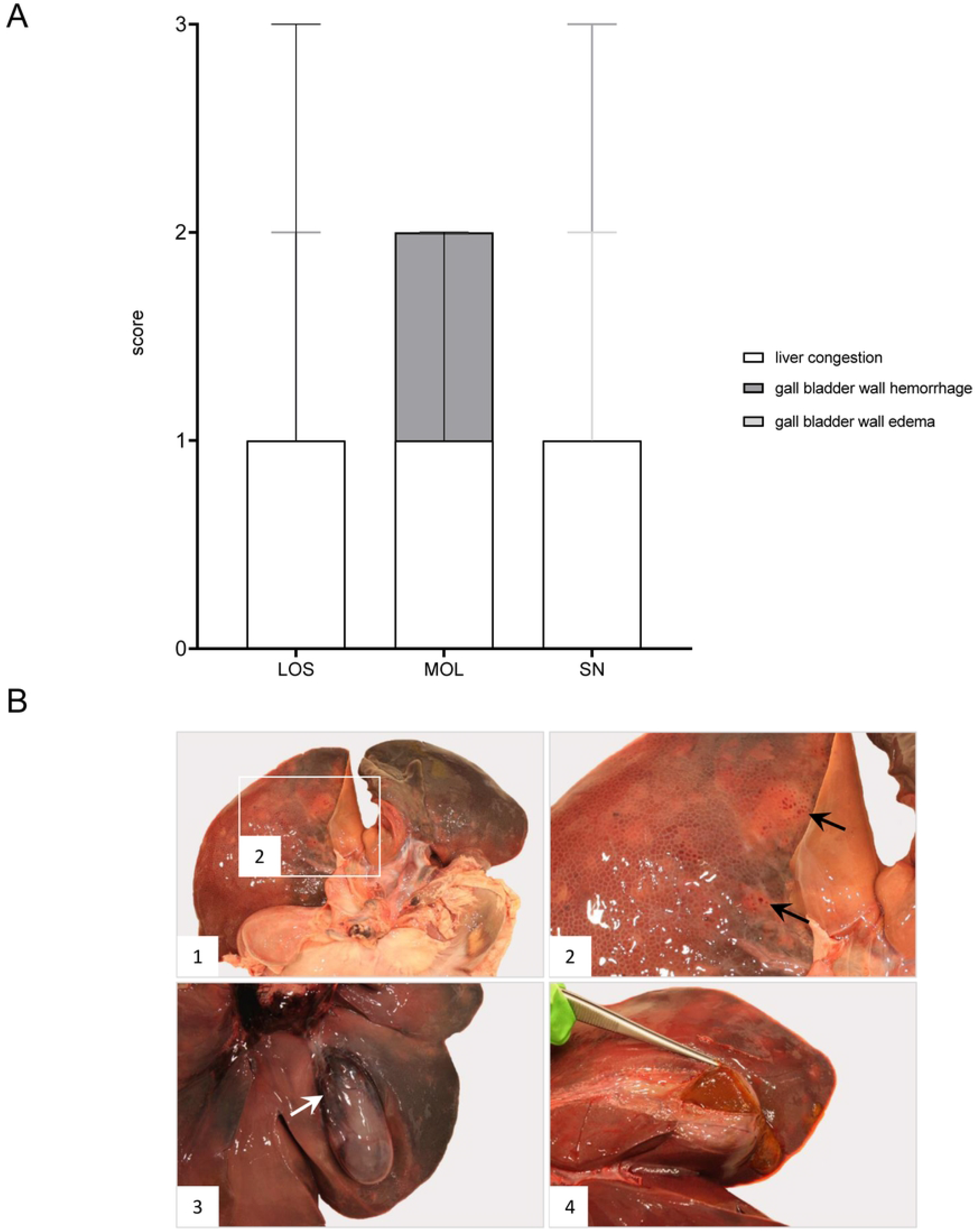
Macroscopical findings of the liver in German ASFV infected wild boar cadavers. A) Stacked bar diagram of gross scoring of hepatic congestion as well as hemorrhage and edema affecting the gall bladder wall on a scale from 0 to 3. Scores are presented as the median value with range. B) Representative macroscopical liver lesions are shown in B1-4. Generally, livers were consistently autolytic to varying degrees (B1-4). In a single wild boar from LOS, multifocal subcapsular pinpoint hemorrhages (arrows) were observed (B1-2). Coalescing hemorrhages of a mildly edematous gall bladder were found in a pig of SN (B3). Viscous biliary sludge was throughout detectable in wild boar from the different outbreak districts (B4).

Due to poor preservation of the liver one animal from LOS had to be excluded from investigation. Congestion of the liver was severely present in two and mildly present in four wild boar. Multifocal subcapsular hemorrhages were observed in one wild boar. While none of the wild boar showed edema of the gall bladder wall, slight hemorrhages were found in a single animal.

Only three livers were evaluated from MOL due to progressive autolytic processes. Mild to moderate congestion was apparent in two livers. Gall bladder lesions could be assessed in 4/5 MOL animals and included mild hemorrhages, but no edematous changes.

All wild boar from SN showed mild congestion of the liver with 1/4 animal revealing a mildly edematous gall bladder wall with moderate hemorrhages.

Wild boar of all districts showed highly viscous bile that was defined as biliary sludge.

##### Histopathology

Histopathological examination was possible only in well-preserved livers (n=5). A summary of histopathological observations is included in Table S2 while representative lesions are illustrated in Fig 16.

**Fig 16.**
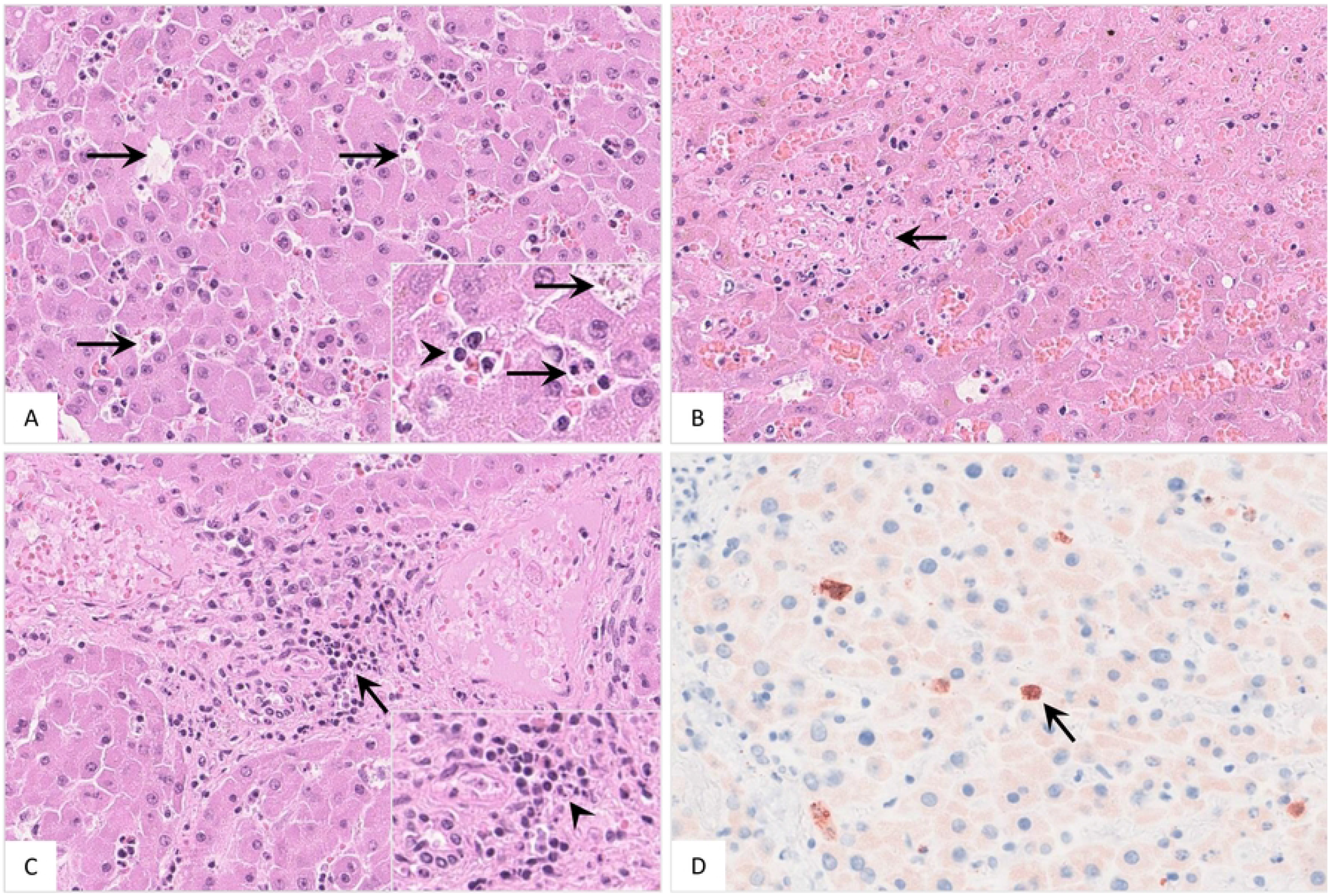
Histopathological results detected in the liver of naturally ASFV infected wild boar from Germany. A) The liver revealed moth-eaten appearance (arrow) due to multifocal apoptosis/necrosis of Kupffer cells (inlay, arrows). The sinusoids were mildly infiltrated by few numbers of neutrophils (inlay, arrowhead). B) In contrast to Kupffer cells, only few hepatocytes showed apoptotic/necrotic changes (arrow). C) There was mild periportal infiltration of mononuclear cells (arrow) which showed cellular degeneration (inlay, arrowhead). D) Mostly moderate numbers of positively labeled Kupffer cells (arrow) were detected in wild boar.

Microscopical lesions included apoptosis/necrosis of Kupffer cells in all available livers (Fig 16A) and to a lesser extent of hepatocytes which was observed in 4/5 wild boar (Fig 16B). Viable and degenerate neutrophils as well as mononuclear cell infiltrates were found in the hepatic sinusoids in four wild boar (Fig 16A) whereas periportal infiltrates consisting of mononuclear cells appeared in all animals (Fig 16C). Autolytic changes aggravated further analysis of the gall bladder. All available livers were analyzed by immunohistochemistry and revealed positive immunolabeling of mainly moderate numbers of cells phenotypically consistent with Kupffer cells (Fig 16D) except for three wild boar from SN which showed rather low antigen levels.

#### Stomach and intestine

##### Gross pathology

Due to progressive autolysis the gastrointestinal tract could be evaluated only in individual animals. Macroscopic findings are indicated in Fig 17 while Table S3 provides detailed scoring results.

**Fig 17.**
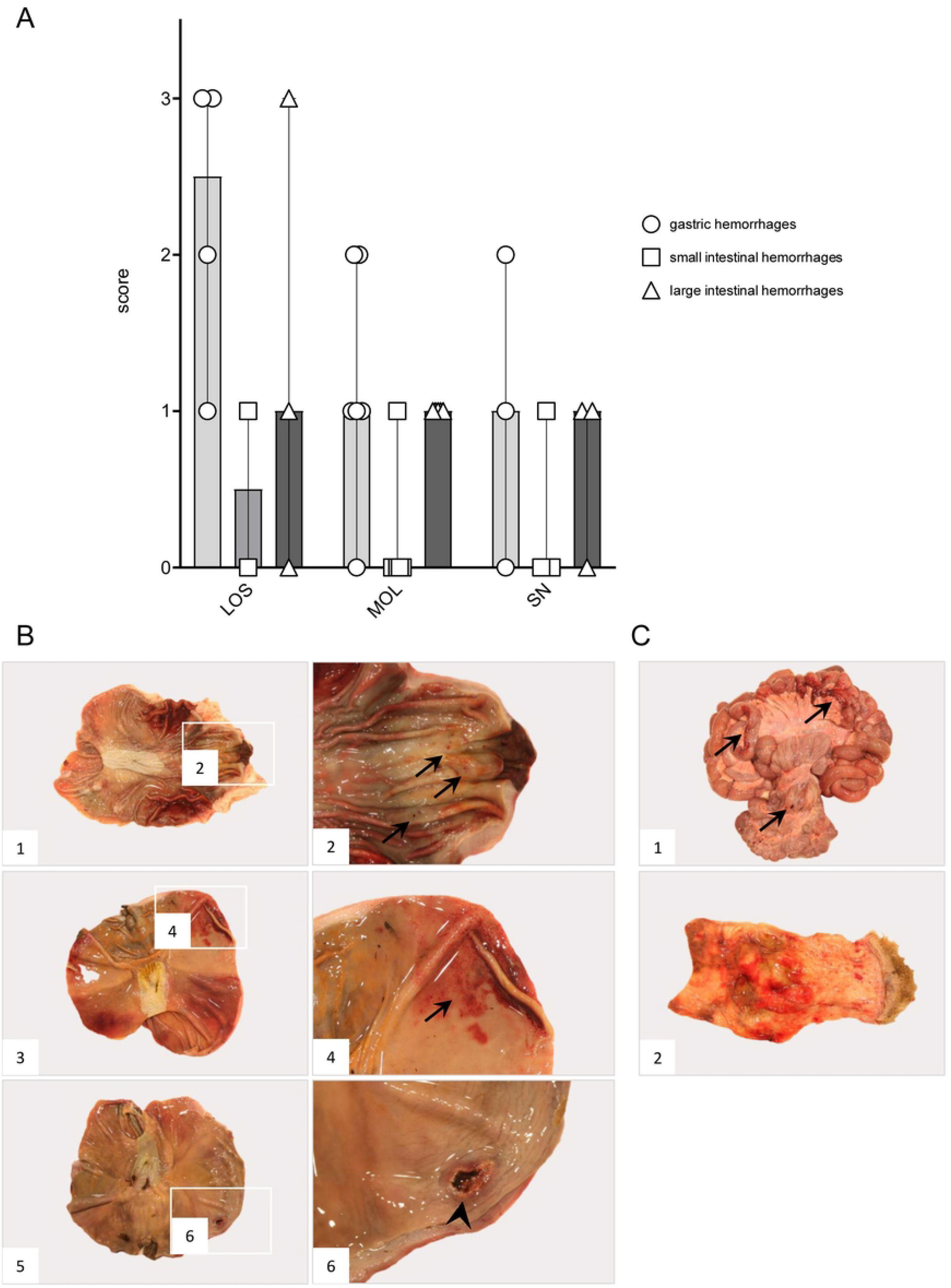
Gross pathology of the gastrointestinal tract in naturally ASF infected wild boar cadavers from German outbreak areas LOS, MOL and SN. A) Stacked bar diagram demonstrating the median values with range of criteria investigated in wild boar. The gastrointestinal tract was investigated for serosal and mucosal hemorrhages, if not already autolytic. Hemorrhages of the stomach, small and large intestine were scored from 0 to 3. B) Pathological alterations in the stomach were found in almost all wild boar which were possible to investigate. Multifocal to diffuse pinpoint hemorrhages (arrows) were found in wild boar from LOS (B1-2) and SN (B3-4). Multiple chronic ulcers of the gastric mucosa (arrowhead) were detected in an animal from the SN district (B5-6). C) In most cases the intestine showed progressive autolysis. While paintbrush hemorrhages (arrows) were sporadically observed on the small and large intestinal serosa (C1), multifocal to diffuse rectal mucosal hemorrhages occurred more frequently (C2).

In 4/7 animals from LOS the stomach could be evaluated while advanced autolysis prohibited investigation of other animals. In these animals, hemorrhagic gastritis of differing severity was detected. The small intestine showed progressive autolysis in five pigs. In one out of the two well preserved intestines, serosal multifocal pinpoint hemorrhages were found. As observed for the small intestine, also the large intestine was severely autolytic. Here, only in 2/7 pigs multifocal to diffuse hemorrhages were found on the serosal and mucosal rectal surface. Hemorrhagic ascites was detected in one animal.

In 4/5 animals from MOL mainly mild hemorrhagic gastritis was present. One animal revealed multifocal hemorrhages in the small intestinal serosa whereas all wild boar had multifocal hemorrhages in the large intestine, especially affecting the rectal mucosa. In 2/5 wild boar hemorrhagic ascites was present.

Due to progressive autolysis of the gastrointestinal tract of only 3/4 animals from SN was investigated. In two wild boar mild or moderate hemorrhagic gastritis, respectively, was present. Chronic ulcerative gastritis was found in one animal. Multifocal serosal hemorrhages were detectable in one wild boar whereas hemorrhages in the large intestine occurred in two animals mainly affecting the rectal mucosa. Two wild boar revealed chronic fibrinous peritonitis.

Histopathological examination was not carried out due to advanced autolysis of the gastrointestinal tract.

### Nervous system

#### Brain

##### Gross pathology

Hemorrhages of the brain were detected in 2/5 animals from MOL in the cerebellum or thalamus, respectively (Fig 18A). Both the cerebellum and cerebrum were further evaluated by histopathology since data on respective lesions are sparse.

**Fig 18.**
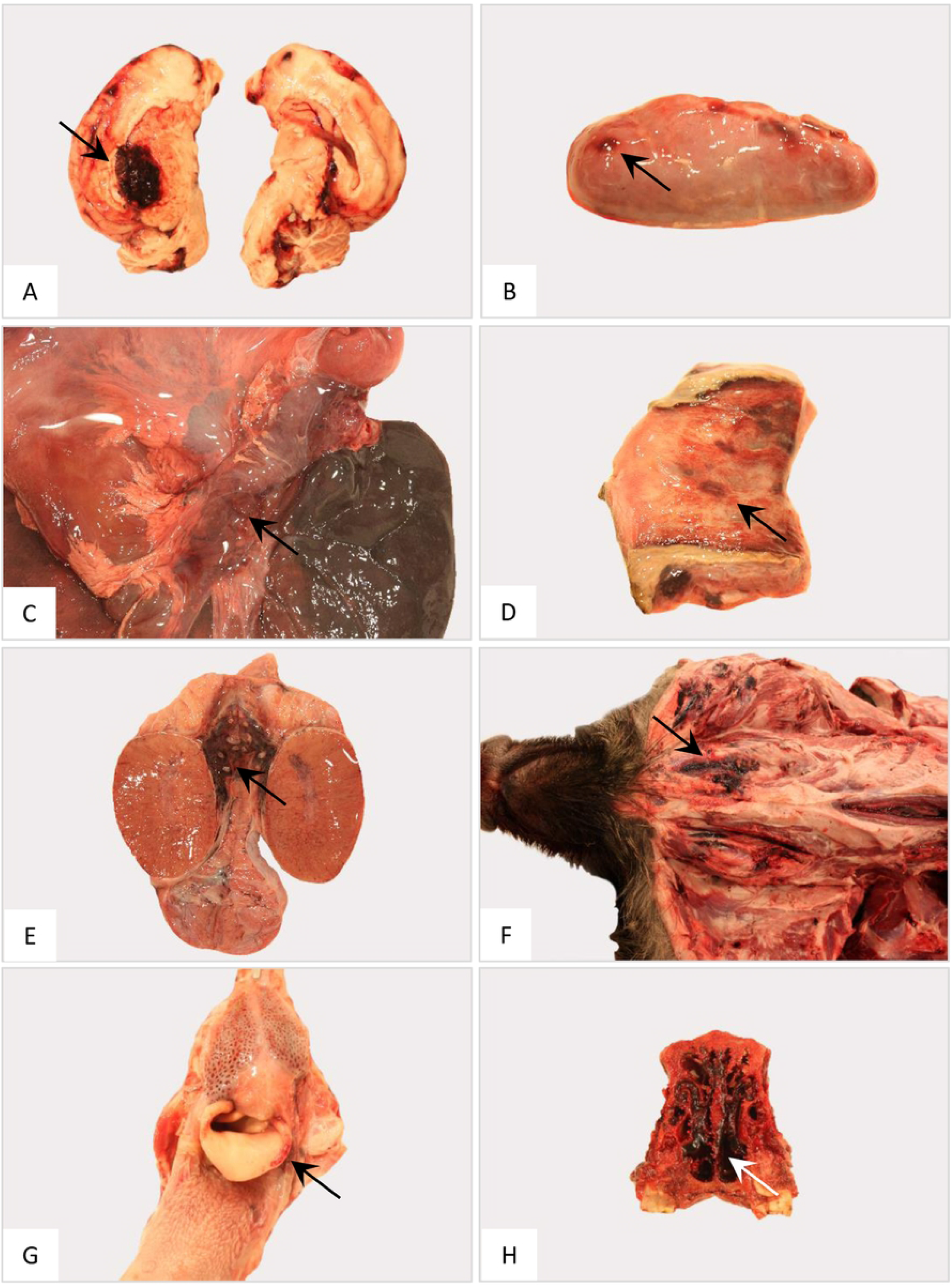
Gross pathology of the nervous, endocrine and reproductive organ systems and other findings in naturally ASFV infected wild boar cadavers from Germany. Representative lesions included hemorrhages in the cerebrum (A), adrenal gland (B), pancreas (C), vestibulum vaginae (D), testis (E), subcutaneous tissue (F), larynx (G) and nasal mucosa (H). Arrows indicate hemorrhagic changes in the respective organs.

##### Histopathology

Brains of all wild boar were evaluated histopathologically as indicated in Table S3. Representative microscopical findings of the cerebellum and cerebrum are depicted in Fig 19 and 20, respectively.

**Fig 19.**
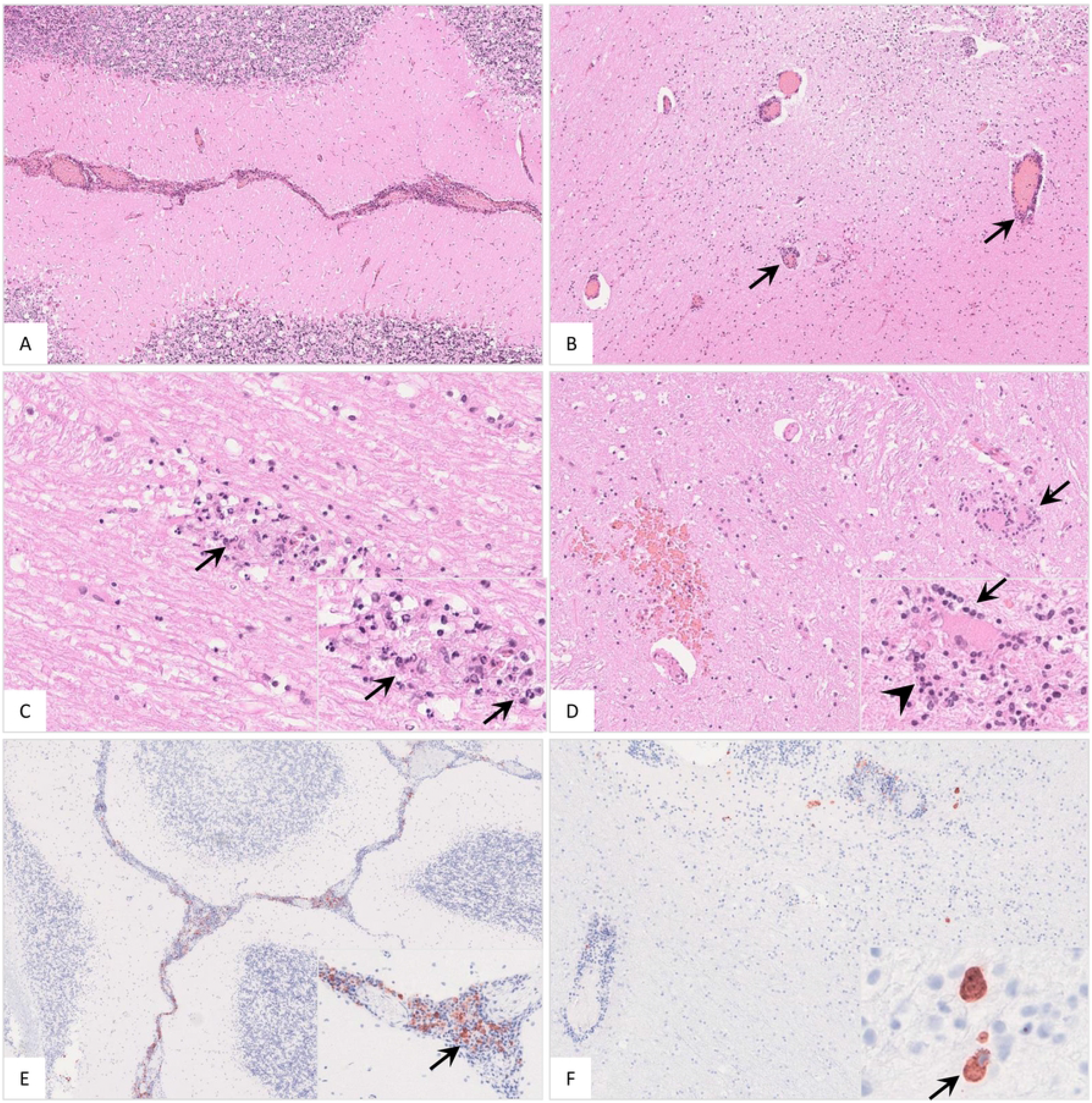
Histopathological findings in the cerebellum of ASFV infected wild boar. A) Meningitis was present in affected animals. B) Cerebellar encephalitis was characterized by multifocal perivascular cuffs consisting of mononuclear cell infiltrates. C) Parenchymal mononuclear infiltrates (arrow) showed multifocal apoptosis/necrosis (inlay, arrow). D) Hemorrhage (left), perineural satellitosis (arrow, also see inlay) and microgliosis (inlay, arrowhead) were recognized. E and F) Cerebellar meninges as well as brain parenchyma revealed positively labeled macrophages of differing amounts (inlays, arrow).

**Fig 20.**
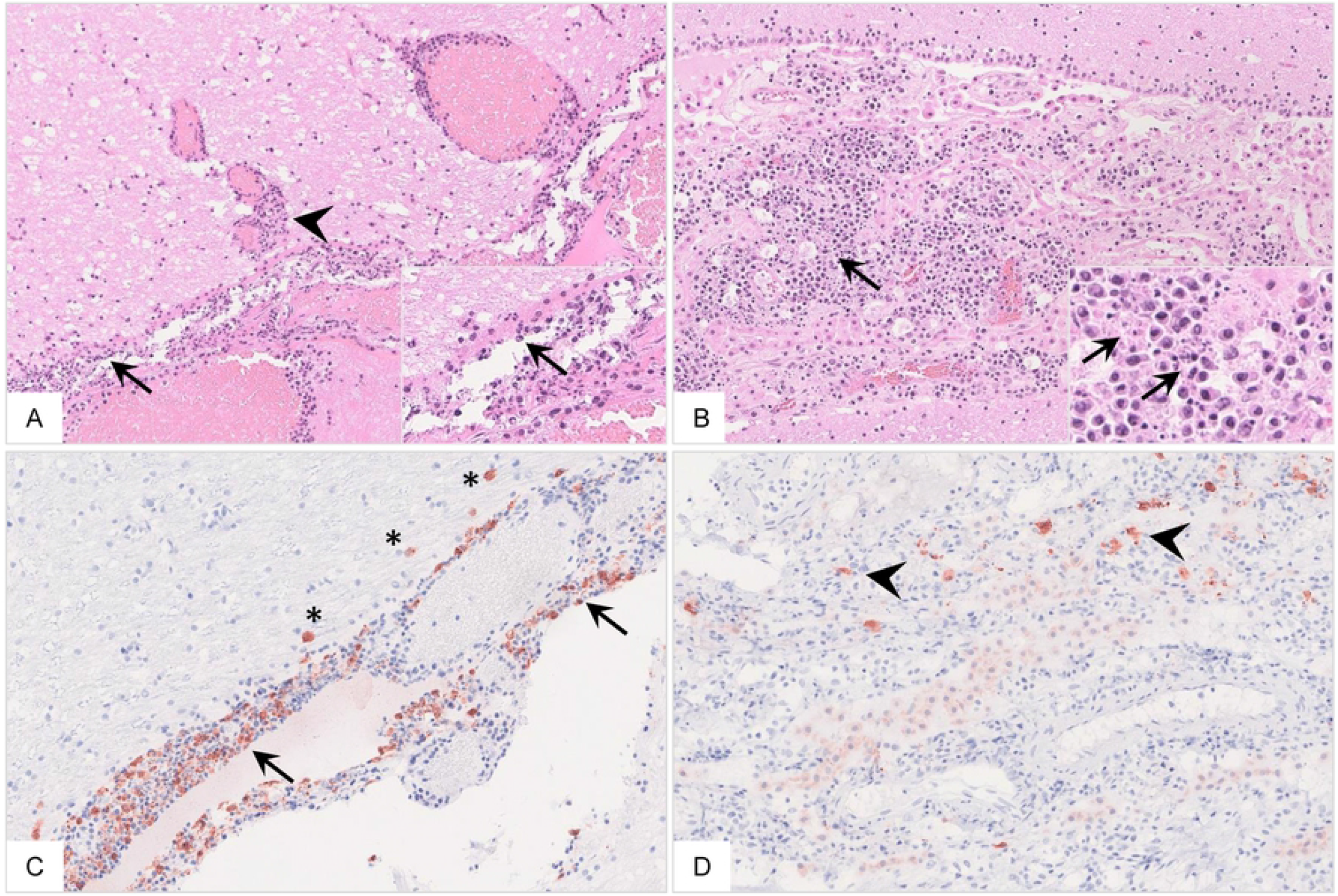
Histopathology of the cerebrum of ASFV infected wild boar carcasses. A) The meninges (arrow) and adjacent brain parenchyma (arrowhead) were infiltrated by mononuclear cells via Virchow Robin spaces. Mononuclear cells showed multifocal apoptosis/necrosis (inlay, arrow). Meningeal vessels were prominently dilated. B) Mononuclear inflammation was limited to the choroid plexus within ventricles (arrow) with multifocal apoptosis/necrosis of infiltrating cells (inlay, arrow). There was degeneration of only few plexus epithelial cells. C and D) Immunopositive cells were present to variable extents in the meninges (arrow), brain parenchyma (asterisk) as well as in the choroid plexus epithelium (arrowhead) phenotypically consistent with macrophages.

In the majority of animals originating from LOS, inflammatory processes were found in both the cerebellum and cerebrum. Inflammation of the cerebellum was found in 3/7 animals while the cerebrum was affected in 6/7 animals. Cerebellar meningitis was present in three animals, two of them had additional encephalitis. Cerebral meningitis was found in five animals while encephalitis was less frequent and occurred only in two wild boar. However, three animals showed inflammation limited to the plexus choroideus. Hemorrhages in the cerebrum were found in two animals.

In contrast to animals from LOS, all pigs from MOL showed cerebellar and/or cerebral inflammation, respectively. Whereas meningitis was detected in all wild boar, encephalitis was diagnosed in four pigs. One wild boar additionally showed cerebellar parenchymal hemorrhage. As the cerebellum, the cerebrum was affected by meningitis in four animals, by encephalitis and by plexus choroiditis in three pigs. Hemorrhagic changes were seen in one wild boar.

In animals found in SN, the cerebellum showed changes in 3/4 animals while the cerebrum was affected in all boar. Only one pig showed cerebellar meningeal inflammation while three animals displayed cerebellar encephalitis. Similar findings were obtained in the cerebrum with meningitis noted in one and plexus choroiditis found in all animals. Neither the cerebellum nor the cerebrum showed any hemorrhagic changes.

Generally, inflammatory infiltrates mainly consisted of large numbers of partly degenerate mononuclear cells including macrophages, lymphocytes and some plasma cells. Immunohistochemical results varied widely among animals, ranging from a low to a large amount of viral antigen-positive cells showing macrophage morphology. In three animals from SN, in one from MOL and in one from LOS the cerebellum was immunohistochemically negative. In the cerebrum, antigen was detected in all pigs except for one animal from LOS.

### Endocrine system

#### Adrenal gland

##### Gross pathology

In the adrenal gland multifocal to coalescing hemorrhages were observed in 3/7 animals from LOS, but also occurred in one animal from SN (Fig 18B). Since data on adrenal histopathology upon ASF infection is missing, the organ was investigated microscopically.

##### Histopathology

In the majority of animals, the adrenal gland was already autolytic and not suitable for histopathological investigation. Nevertheless, the adrenal glands of 3/7 animals from LOS, 1/5 animals from MOL, and 2/4 animals from SN could be examined microscopically. Individual histopathological results are listed in Table S3 and are visualized in Fig 21.

**Fig 21.**
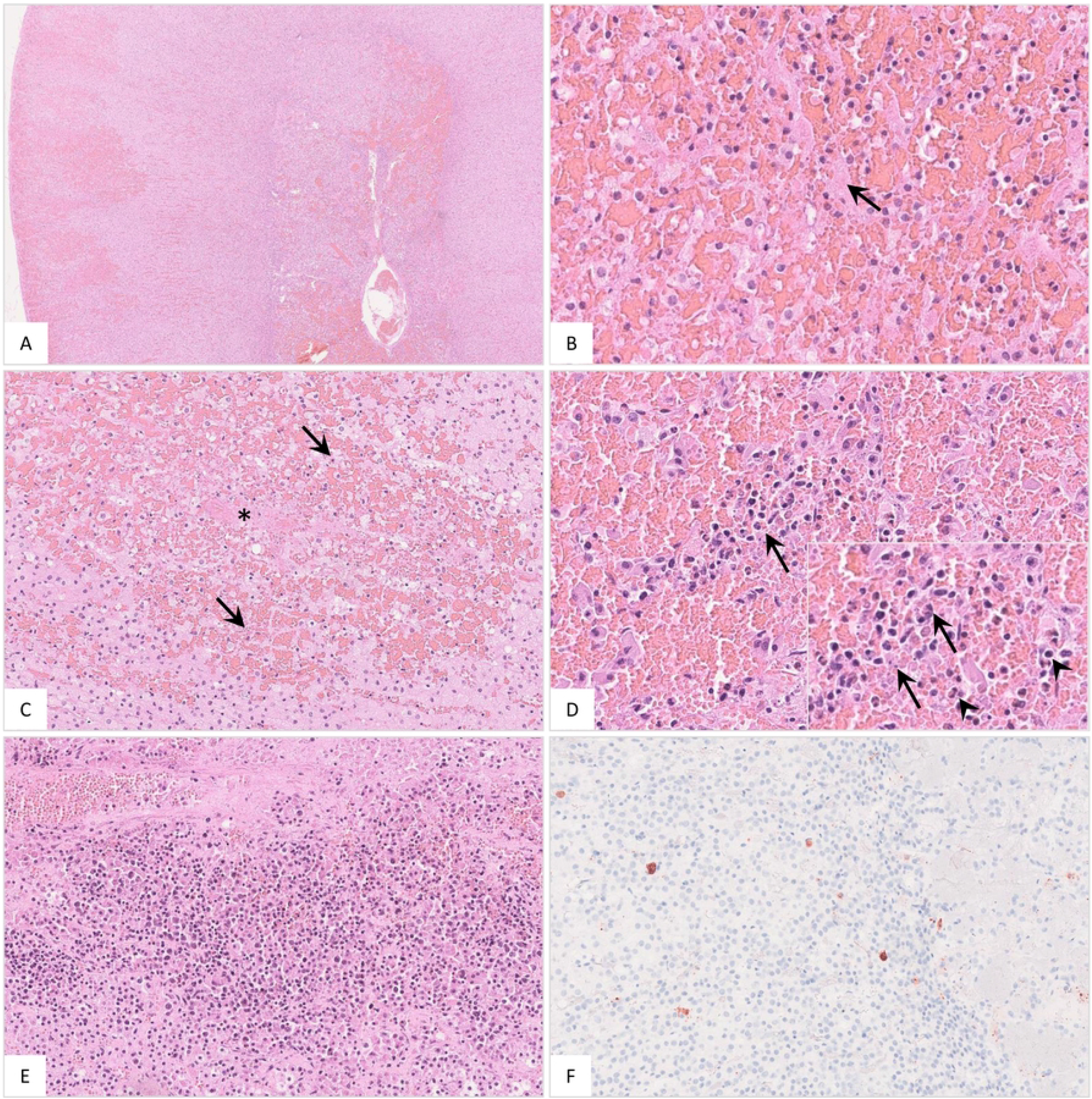
Histopathological findings of the adrenal gland in ASFV infected wild boar. A) Overview of the adrenal gland of a deceased wild boar. The adrenal gland showed extensive cortical and medullary hemorrhages. B) Multifocally, fibrin thrombi were visible in the sinusoids (arrow). C) Occasionally, areas of necrosis were present in the cortex (arrow). There was fibrin deposition (asterisk) and massive hemorrhage in the affected location. D) The medulla was markedly expanded by hemorrhage. Infiltrating mononuclear cells as well as few neutrophilic granulocytes (inlay, arrowhead) accumulated around degenerated cells (inlay, arrow). E) The adrenal medulla was severely infiltrated by mononuclear cells admixed with fewer neutrophils. F) Moderate amounts of antigen-positive macrophages were found in the majority of animals.

Pathological changes of the adrenal gland were observed in three animals from LOS. Hemorrhages could be found affecting the cortex and medulla. In addition, sinusoidal thrombosis and single necrotic areas, either in the cortex and/or medulla, were present. Inflammation of the adrenal medulla and to a lesser extent of the cortex were diagnosed in all three wild boar from LOS. The inflammatory infiltrate mainly consisted of viable and degenerate mononuclear cells admixed with few neutrophils. Similar findings were observed in the affected animals from MOL and SN. Immunohistochemistry revealed moderate amounts of antigen in cells morphologically consistent with macrophages.

Moreover, in wild boar from SN, severe diffuse pancreatic edema and hemorrhage were detected in 2/4 animals (Fig 18C).

### Reproductive system

Multifocal foci of hemorrhage were found in the vaginal vestibulum (Fig 18D) in one wild boar from LOS while one animal from MOL revealed diffuse hemorrhages and edema in the spermatic cord (Fig 18E).

### Other findings

Subcutaneous hemorrhages were present in two animals from LOS (Fig 18F). Furthermore, multifocal hemorrhages in the epiglottis (Fig 18G) as well as epistaxis (Fig 18H) could be detected in one wild boar each from SN.

Furthermore, verminous pneumonia was present in all wild boar from LOS and in one animal each from MOL and SN. One animal from SN showed multiple chronic gastric ulcers. However, the latter lesions were not included in pathological scoring since they cannot be specifically attributed to ASF.

Bacteriologic investigations of secondary bacterial infections were not performed for biosecurity reasons.

### Antibody detection against African Swine Fever Virus

All animals were tested for anti-ASF antibodies by IPT as shown in S1 Fig. Except for one animal from LOS, all wild boar developed antibodies of different titers between 200 and 800. Higher titers tended to be found in the animals from MOL, titers ranging from 200 to 1600, and in two wild boar from SN having titers of 800 and 3200, respectively. One animal from SN showed a titer of 40. In the fourth wild boar from SN no test could be performed due to limited sample material.

For comparison, three domestic pigs from a previous study inoculated with the moderately virulent ASFV strain “Estonia 2014” were analyzed for anti-ASFV-specific antibodies. Starting at day 14 pi all pigs developed antibody titers between 200 and 400. Since one pig had died at day 14 pi, only two animals could be analyzed in the following days. On day 21 pi, titers increased to 800 and 1600. On day 28 pi titers further increased to 3200 or even remained at the same level of 1600 while on day 35 pi antibodies dropped in one animal to 800, but increased in the other pig to 3200. On day 41 pi a second increase of the titer to 1600 was noted in one pig whereas in the other one, antibodies remained constantly high at 3600.

**Suppl. Figure 1. Antibody titers determined by immunoperoxidase test in German wild boar cadavers compared to ASFV “Estonia 2014” experimentally infected domestic pigs on different days pi.** LOS = Landkreis Oder-Spree, MOL= Märkisch-Oderland, SN= Spree-Neiße, EST= Estonia 2014

## Discussion

Filling the documentation gap on the pathology after ASF field infection, the aim of the present study was to examine ASFV infected wild boar that succumbed to the disease under natural conditions in both virological and pathomorphological detail. Furthermore, the impact on the virulence of emerging virus variants II, III and IV in ASF outbreak areas of Eastern Germany was analyzed.

A total of 16 wild boar aged between 0 and 2 years of different sexes were investigated. Despite the different preservation status, the organs of each animal could be examined for ASFV genome load and revealed consistently positive results. Significant differences were not found between outbreak areas and the different variants, but wild boar from SN tended to show both lower viral genome loads and viral antigen scores compared to animals from LOS and MOL. However, especially the viral genome load has limited informative value at this point since viral genome can be detected up to 100 days after infection (28) and the time at which the genome load decreases varies greatly between experiments (11, 18, 19).

In addition to organ-wide detection of viral genome, all wild boar irrespective of the outbreak area and virus variant were diagnosed with classic and severe ASF lesions resembling a systemic hemorrhagic disease (6). While some unique macroscopic findings have not been described so far, many of the lesions found in the present wild boar study have been shown already experimentally (29), but however, experimental findings do not reflect the severity and diversity seen under field conditions. Nevertheless, striking, but not significant differences were evident between the variant groups. Interestingly, the highest total score for gross pathological changes was given for wild boar from SN infected with variant IV, followed by animals from MOL infected with variant III which showed an intermediated total score and wild boar from LOS infected with variant II having the lowest macroscopical score.

For ASF, four different courses of the disease have been described and include peracute, acute, subacute, and chronic stages, which are associated with typical lesions (6). Petrov et al. (28) moreover specified the subacute stage as chronic-like and differentiated into lethal and transient course after infection with moderately virulent ASFV. Gross pathomorphological changes of the subacute/chronic-like stage include multifocal hemorrhages, edema, lymphadenitis, interstitial pneumonia and ascites (6, 28) whereas bacterial secondary infections inducing fibrinous polyserositis, chronic pneumonia and necrosis of tonsils, however without vascular changes, predominate in chronic courses (6). Lesions in acutely and chronically ASFV infected domestic pigs were also already presented in detail decades ago (12). The animals with chronic disease showed comparable lesions as observed in the acutely infected pigs, but additionally revealed chronic changes particularly including pericarditis, pneumonia and lymphadenitis. In the present study, in contrast to the animals from LOS and MOL, wild boar from SN more frequently showed lesions indicative for a lethal subacute protracted disease course.

More specifically, chronic inflammatory processes such as fibrinous pericarditis, pleuropneumonia and peritonitis were more frequently detected in SN animals. At the same time, wild boar from SN, and to a lesser extent also animals from MOL showed more severe hemorrhages in the urinary bladder and bone marrow, but fewer acute hemorrhages as detected in the hearts of animals from LOS. In wild boar experiments with detailed pathomorphological investigation, only mild, pinpoint hemorrhages of the urinary bladder, variable hemorrhages of the heart and congestion of the bone marrow have been mainly observed in acutely infected animals that succumbed to highly virulent infection with ASFV “Armenia07” while extensive hemorrhages in these organs were rather absent (29).

Based on this, and in line with virological and immunohistochemical data, this may indicate that at least wild boar infected with the SN variant experienced a more protracted disease course than pigs from LOS suggesting a slightly decreased virulence of the SN virus variant IV to wild boar that still led to the death of the respective animals. However, considering the small number of cadavers and the indefinite sample material, this should be interpreted with caution and must be confirmed experimentally under standardized conditions in any case.

Although the majority of organs could be assessed macroscopically, we had to refrain from a detailed semiquantitative histopathological analysis because autolysis had already progressed too far in some cases, which would have considerably reduced the number of samples for investigation. However, in line with macroscopic findings, histopathology confirmed the severe course of disease in all animals regardless of the outbreak area and the virus variant. Since most wild boar studies focus only on macroscopic pathology, it is even more important to study the histopathology of natural ASF infection in more depth (17, 18, 29, 30).

For example, adrenal hemorrhages, which have been described to occur in wild boar upon experimental infection (29), were examined in more histopathological detail and revealed interesting results in the present study. Our findings mirror a condition known as Waterhouse Friderichsen syndrome (31). It has been correlated with several bacterial and viral diseases and is characterized by severe hemorrhage, necrosis and microvascular thrombosis. Although the pathophysiology is not fully understood hemorrhages are explained by stress-induced release of adrenaline, vasculitis and coagulation disorders including disseminated intravascular coagulation. In line with the latter, microvascular thrombosis could be shown in multiple organs as signs of acute organ injury of naturally infected wild boar in this study (12).

Of note, histopathology further highlighted the unique finding of localized inflammation of the cerebral choroid plexus, which occurred in wild boar irrespective of the outbreak district, but mainly affected the majority of animals from MOL and SN. So far there are only minor reports on ASF lesions in the central nervous system (12, 32) which can occur at all stages of the disease as demonstrated by Moulton and Coggins (12) in acutely and chronically succumbing as well as in surviving pigs. In addition to mononuclear infiltration of meningeal and cerebral vessels, perivascular hemorrhage, occasional vascular thrombosis and neuronal degeneration, necrosis of the choroid plexus epithelium has been described once in few acutely infected animals (12). Naturally infected wild boar presented in this study showed pronounced mononuclear inflammation with massive cell deaths in addition to occasional necrosis of the plexus epithelium, again suggesting a longer disease course at least in animals from SN.

To further extrapolate how long naturally infected wild boar might have lived with the disease, antibody titers were determined and compared to those of surviving ASFV “Estonia 2014” experimentally infected domestic pigs from a previous trial. In domestic pigs, low antibody titers were detectable from day 14 to a maximum titer of 400, then increased to a max of 3200 by day 28, and remained constantly high until 41 days post infection, at least in one domestic pig. However, the other pig showed a drop from 3200 to 800 on day 35 pi and a second subsequent increase. While it cannot be excluded that a consumption or decay of antibodies occurred, one should also consider measurement inaccuracies of the semi-quantitative test when targeting the fluctuant antibody titers. When comparing this to wild boar which showed titers of at least 200, the majority of animals independent of the outbreak area might have lived with ASF for more than 14 days.

As suspected based on the pathological data in animals from SN, but also MOL, the course of the disease was probably longer since they tended to show higher titers of max 3200 and 1600, respectively, while wild boar from LOS reached titers of only max 800. Surprisingly, antibody titers showed no clear correlation to the chronicity of lesions observed in several wild boar since one animal from SN with chronic lesions produced only minimal antibody titers. On the one hand, the chronic lesions in this animal could have already existed before and might not necessarily associated with ASFV infection. On the other hand, as hypothesized above, antibodies may have declined over time. To date, little is known about the host’s immune response against ASFV, but it is of general acceptance that antibodies directed against ASFV are not sufficient for protection against the disease (33). However, experiments to investigate the dynamics of antibody development in ASF could be useful to draw conclusions on the disease in wildlife.

## Conclusion

In summary, this is the first study describing the lesion spectrum in wild boar succumbing to ASF after infection with different virus variants that have emerged within one year in Germany. Virological and pathomorphological data suggest possible differences in the virulence of the variants. At least wild boar infected with the SN variant IV experienced a more protracted but nevertheless lethal disease course compared to animals infected with LOS variant II or the MOL variant III which is more likely to be classified as intermediate. These findings are particularly important with regard to the spread and continued occurrence of ASFV in endemic areas. To elucidate the pathogenicity and differences in the virulence and disease dynamics of the emerging virus variants more thoroughly, further experimental studies in wild boar as well as comparative investigations in domestic pigs under late human endpoint conditions are urgently needed. These studies should also address the impact of protracted disease courses on shedding and thus transmission characteristics.

## Acknowledgement

We would like to thank the Brandenburg Crisis Center and the veterinary authorities in the districts Märkisch-Oderland, Oder-Spree and Spree-Neiße for their help and the submission of the examined wild boar carcasses. Further we would like to thank Christian Loth, Silvia Schuparis, Ulrike Kleinert and Robin Brandt for their valuable support during necropsies and in the laboratory.

## Supplementary Material

**Table S1, Summary of individual organ genome copy numbers in wild boar**

**Table S2, Summary list of histopathological changes and immunohistochemistry results in wild boar**

**Table S3, Summary list of gross lesions scored on a semiquantitative scale in wild boar**

**Figure S1, Antibody titers determined by immunoperoxidase test in German wild boar cadavers compared to experimentally infected domestic pigs on different days pi.**

## Author contributions

Conceptualization, J.SE. and S.B.; Methodology, J.SE., S.B., P.D., A.B.; Validation, J.SE., S.B.; Formal Analysis, J.SE., P.D.; Investigation, J.SE., S.B., P.D., A.B.; Resources, S.B. and J.SE.; Data Curation, J.SE., P.D.; Writing – Original Draft Preparation, J.SE., P.D.; Writing – Review & Editing, J.SE., S.B., P.D., A.B.; Visualization, J.SE., P.D.; Supervision, J.SE. and S.B.

## Informed Consent Statement

Not applicable.

## Data Availability Statement

All data available can be obtained on request from the corresponding author.

